# Potential of exogenous biological nitrification inhibitor addition to improve soil nitrogen availability for crop growth

**DOI:** 10.64898/2026.07.11.738001

**Authors:** Paula A. Rojas-Pinzon, Bernhard Seidl, Stella Kejik, Iris Karbon, Christopher J. Sedlacek, Judith Prommer, Katrin Pilz, Christoph Bueschl, Taru Sandén, Heide Spiegel, Andrew T. Giguere, Petra Pjevac, Lucia Fuchslueger

**Affiliations:** University of Vienna, Centre for Microbiology and Environmental Systems Science, Department for Microbiology and Ecosystem Science, Vienna, Austria; University of Vienna, Doctoral School in Microbiology and Environmental Science Vienna, Austria; Department of Agrobiotechnology IFA-Tulln, Institute of Bioanalytics and Agro-Metabolomics, University of Natural Resources and Life Sciences, Vienna, Austria; University of Southern Indiana, Department of Biology, Evansville, IN, USA; Austrian Agency for Health and Food Safety (AGES), Department for Soil Health and Plant Nutrition, Vienna, Austria; Joint Microbiome Facility of the Medical University of Vienna and the University of Vienna, Vienna, Austria; Environment and Climate Hub, University of Vienna, Austria

**Keywords:** Biological nitrification inhibitor, nitrogen immobilization, mineralization, MBOA, DMPP, MHPA, Limonene

## Abstract

Modern agriculture is characterized by substantial fertilizer nitrogen (N) losses from soils, resulting in low crop N-use efficiency. Biological nitrification inhibitors (BNIs) are studied as a strategy to improve N retention in soils by suppressing nitrification. However, the impacts of applying exogenous BNIs to crops with unknown intrinsic BNI capacity remain poorly understood. In this study, we evaluated the impacts of adding three BNIs (methyl 3-(4-hydroxyphenyl) acrylate [MHPA], 6-methoxy-2(3H)-benzoxazolone [MBOA], and limonene), their mixture, and the synthetic nitrification inhibitor 3,4-dimethylpyrazole phosphate (DMPP) on barley (*Hordeum vulgare* L.) growth, plant and soil N dynamics, and soil microbial communities. Using a rhizobox system with planted and bare-soil compartments, combined with ^15^N isotope tracing and molecular microbial community analyses, we assessed the spatio-temporal dynamics of N transformations, losses, plant N uptake, and microbial community responses in an alkaline agricultural soil. Independent of inhibitor application, the applied fertilizer N was lost primarily through NO₃⁻ leaching (3-9% of the applied N). In contrast, N₂O emissions represented only 0.001–0.028% of the applied N and varied with inhibitor type. MHPA increased dissolved inorganic N soil pools without affecting plant biomass or ^15^N uptake or strongly shifting microbial community composition. MBOA reduced NO_3_^-^ concentrations in soil pore water without influencing plant growth or N uptake but shifted soil microbial community composition. In contrast, limonene reduced plant growth and ^15^N uptake and most significantly altered microbial community composition, without significantly changing N availability. Applying a BNI mixture, as well as limonene alone, was detrimental to plant growth and ^15^N uptake. DMPP showed only minor effects on N pools, plant growth, plant N uptake and microbial community composition. Overall, our results reveal both the potential and limitations of exogenous BNI application for improving N retention in crop systems.

## Introduction

To sustain the food supply for a current global population of approximately 8 billion people, agriculture heavily relies on synthetic nitrogen (N) fertilizer, which accounts for about 60% of total N inputs to the environment (FAO, 2025). Although the extensive use of N fertilizer has increased crop yields, estimates of crop N use efficiency (NUE) have not improved much in the last 60 years, remaining between 40 and 56% (FAO, 2025). As a result, approximately half of the fertilizer N is lost, leading to significant environmental damage. Losses vary widely depending on climate, soil properties, crop type and management practices, but it is estimated that globally between 10-20% of the N applied is lost through ammonia (NH_3_) volatilization, 10-30% through nitrate (NO_3_^-^) leaching, and about ∼1% as nitrous oxide (N_2_O) emissions (Tian et al., 2020; Ma et al., 2021; Wang et al., 2025). Under the current conditions, reducing N losses is a critical first step toward achieving a sustainable agriculture that can reliably meet the food demands of a growing global population.

A central process in the N cycle, directly linked to N fertilizer losses, is nitrification, the microbially mediated oxidation of ammonia (NH_3_) to NO_3_^-^. Nitrate, like ammonium (NH_4_^+^), represents a readily available form of N for plant uptake (Gigon and Rorison, 1972; Britto and Kronzucker, 2013). However, NO_3_^-^ is highly mobile in soils and serves as a substrate for denitrification. Thus, nitrification can directly and indirectly increase N fertilizer losses (Subbarao et al., 2013; Norton and Ouyang, 2019). Consequently, controlling nitrification activity has emerged as a way to reduce N losses and increase soil N availability for plant uptake (McCarty and Bremner, 1986; Rodgers, 1986; Subbarao et al., 2006). Synthetic nitrification inhibitors (SNIs), such as 3,4-dimethylpyrazole phosphate (DMPP), have shown high efficacy in reducing NO_3_^-^accumulation under certain conditions and in certain soil types (Shi et al., 2016; Zhou et al., 2020; Bachtsevani et al., 2021; He et al., 2023). However, the variable efficacy, high costs, and potential off-target effects of SNIs have shifted the attention towards the discovery of naturally derived inhibitory compounds (Subbarao et al., 2012, 2013; Coskun et al., 2017).

Biological nitrification inhibitors (BNIs) are compounds produced by some plants as an adaptation to compete for N in oligotrophic environments, as well as in highly fertilized soils where N is prone to loss due to microbial activity (Lodhi and Killingbeck, 1980; Subbarao et al., 2012; Lata et al., 2022). Over the past 40 years, several BNIs have been identified in both perennial and annual plants, yet the BNI capacity of some economically relevant crops, such as barley, is poorly understood (Subbarao et al., 2015; Nardi et al., 2020; Lata et al., 2022). Once a potential BNI compound has been isolated and identified, the next step is typically to test its inhibitory activity in pure cultures or in soil incubations (Nardi et al., 2013; Sun et al., 2016; Otaka et al., 2021; Kaur-Bhambra et al., 2022; Kolovou et al., 2023). However, less studies have examined the effects of adding exogenous BNIs to plant-soil-microbes systems (Yao et al., 2020; Lan et al., 2022). Such experiments, however, are crucial to properly assess BNI applicability. For instance, compounds such as limonene and other terpenoids, known for their nitrification-inhibiting activity (White, 1988, 1991) also exhibit allelopathic effects (Zhao et al., 2009; Li et al., 2020). This suggests that their impact on plant growth at the concentrations at which nitrification inhibition occurs, should also be considered when assessing BNI suitability. Additionally, BNIs can have off-target effects on other soil processes, which may subsequently influence soil biogeochemical cycles and plant health (Ma et al., 2021; Wu et al., 2026). While BNIs have been extensively studied in plants that naturally produce them, the impacts of BNI addition to crops without known BNI activity remain largely unexplored. In particular, it is unclear how exogenous BNIs affect plant biomass, plant N allocation, soil N pools, and the composition of soil microbial communities.

Plant productivity and ecosystem functioning are sustained through complex interactions among plant roots, soil microbial communities, and soil properties (Philippot et al., 2013, 2023; Moreau et al., 2019). Hence, evaluating the impacts of BNI addition in integrated platn-soil systems is needed before we can integrate the use of BNIs as a management practice in agricultural systems. The objective of this study was to evaluate the effects of three individual BNIs; methyl 3-(4-hydroxyphenyl) acrylate (MHPA), 6-methoxy-2(3H)-benzoxazolone (MBOA), and limonene; their mixture, and a synthetic nitrification inhibitor (DMPP) on barley growth. We used a compartmentalised rootbox system, in which planted and unplanted compartments allowed the assessment of both plant and soil responses to exogenous BNI application. This holistic evaluation of BNI application provides crucially needed insight into their suitability as a management strategy in crops lacking inherent BNI potential.

## Methods

### Soil description and sampling

The soil used in the rhizoboxes has previously displayed high nitrification potential (Rojas-Pinzon et al., 2024), thus representing an agricultural system in which NI application would be potentially highly beneficial. This alkaline soil was collected in spring 2024 from a long-term fertilization experiment managed by the Austrian Agency for Health and Food Safety (AGES) (Lehtinen et al., 2014; Spiegel et al., 2018). This long-term fertilization experiment is located in Lower Austria, in the Marchfeld region (48°12′57.2″N 16°37′06.1″E) and the soil is classified as Calcaric Phaeozem (sandy loam: 30.3% sand, 45.6% silt, and 24.2% clay). With a pH in water of 8.5, this alkaline soil had an initial NH_4_^+^ content of 0.39 ± 0.1 µg N g^-1^ dry weight (dw) soil, a NO_3_^-^ content of 15,19 ± 0.8 µg N g^-1^ soil, and a water content of 11.3% (a further characterisation of the soil can be found in Rojas-Pinzon et al., 2024).

The long-term experiment has consisted of constant N and phosphorus fertilization (120 kg N ha^−1^ yr ^−1^, 75 kg P_2_O_5_ ha^−1^ yr ^−1^) and a crop rotation system of 53%–55% cereals and 45%–47% root crops (Lehtinen et al., 2014; Spiegel et al., 2018). At the time of sampling, the site was uncropped and 40 kg from the top soil were collected by manual shovelling from four field replicate plots. The soil was transported to the laboratory, sieved (4 mm mesh size), and stored at room temperature (20°C).

### Root box system assembly

Acrylic rhizoboxes (15 x 20 x 5 cm, Fig. S1) were used to test the effect of exogenous BNIs on the growth and N uptake of summer barley (*Hordeum vulgare* L.). Each rhizobox was assembled with dark panels, except for a transparent front panel. The bottom panel contained drainage holes arranged in three rows of 0.5-mm-diameter holes spaced 1 cm apart. The back panel had a 5 × 5 mm grid of holes (1 mm inner diameter), which were used for fertilizer and NI solution injection. Each of the two side panels contained two holes (4.5 mm inner diameter) located 8 cm and 11 cm from the top, through which rhizons were inserted for pore water collection. The boxes were divided in two compartments: a planted compartment (15 × 12 × 5 cm) and a bare-soil compartment (15 × 8 × 5 cm) by a solid acrylic strip. Beneath each compartment, a tray was placed for leachate collection.

After adjusting soil water content to 20%, the soil was pre-weighed in plastic bags and then packed into the rhizoboxes (i.e. 960 g and 640 g of soil into the planted and bare-soil compartments, respectively) to achieve a bulk density of 1.4 g cm^-3^ in each compartment. The N fertilizer application rate was 117 μg (NH_4_)_2_SO_4_–N g^-1^ dry weight (dw) soil, split into two applications: 40% was mixed with the pre-weighed soil before filling each compartment, and the remaining 60% was injected as a solution of labeled (^15^NH_4_)_2_SO_4_ (0.6M) six days before the final harvest. The treatments consisted of a control without nitrification inhibitor, and five different NI treatments: DMPP, limonene, MBOA, MHPA and a mixture of the three BNIs. DMPP was applied at a rate of 1% of the applied N (1.1 μg g^-1^ dw soil) as a solution injected into the back of the box. To ensure an even distribution, the solution was split into volume-adjusted injections for the planted and bare-soil compartment. The same approach was used to apply limonene at a rate of 1.8 mg g^-1^ dw soil. MBOA and MHPA were applied at rates of 290 and 387 μg g^-1^ dw soil, respectively. Due to their low water solubility, these two BNIs were premixed with the soil before it was added to the planted and bare soil compartments. The BNI mixture consisted of the three BNIs at the concentrations used in the individual treatments. To facilitate the drainage, 150 g and 100g of glass beads (10 mm diameter) were placed at the bottom of the planted and bare soil compartments, respectively.

Summer barley seedlings, pre-germinated in perlite for one week (21°C with 50% relative humidity during the day and 60% at night), were transferred to the planted compartments (4 seedlings per compartment). For each treatment a set of four rhizoboxes were used. Plants were grown for 15 days, as previous studies on plants with known BNI capacity indicate higher BNI exudation in early, pre-tillering growth stages (Jáuregui et al., 2023).

### Irrigation events, pore water, and leachate sampling

To quantify N losses through leaching and to evaluate the efficacy of the NIs in reducing such losses, four irrigation events were conducted over the course of the experiment starting 24 h after rhizobox setup, and on days 5, 9, and 14. During the first irrigation, 60 mL and 40 mL of water were added to the planted and bare-soil compartments, respectively. In the subsequent three irrigation events, the planted compartments received 100 mL of water, while the bare soil compartments received 70 mL. To determine mineral N concentrations in soil pore water, rhizon MOM samplers (5 cm membrane length, 10 cm glass fiber strengthener, 0.15 μm pore size) were placed at depths of 8 and 11 cm in each compartment, ensuring that the porous membrane was in contact with the soil.

After each irrigation event, the rhizon samplers were activated to extract soil pore water by applying suction through a 12 mL syringe connected to each sampler and withdrawing the syringe plunger. To prevent backflow of extracted water into the soil, 7 cm wooden retainers were used to maintain constant suction, ensuring continuous pore water extraction. After 7 h, the volumes of both the leachates and the pore water were recorded, and samples were collected. The leachate samples were first centrifuged (10 min at 13,751 *g*) and then, together with the pore water samples, stored at-20 °C until analysis. NO_2_^-^ and NO_3_^-^ concentrations in the leachates and pore water samples were quantified with the acidic VCl_3_/Griess reaction (Miranda et al., 2001), while a modified photometric indophenol reaction method (Mulvaney, 1996) was used to quantify NH_4_^+^ concentrations. Cumulative N losses from the leachates were calculated by adding the total mineral N concentrations (µg NO_2_^-^, NO_3_^-^ and NH_4_^+^ g^-1^ dw soil) in each treatment within each compartment over the four sampling points.

### Gas flux measurements

To estimate the N losses as nitrous oxide (N_2_O) emissions, N_2_O concentrations were measured following each irrigation event using static plastic chambers installed in each compartment. The chambers were inserted into the soil to enclose the upper 3 cm, leaving a headspace volume of 37.2 mL. Each chamber was sealed with a screw cap fitted with a butyl septum, through which gas samples were collected. Chambers were closed immediately after irrigation, remained closed during the first 24 h, and were left open until the next irrigation event. An initial gas sample (12 mL) was taken after the chamber was closed and transferred to 15 mL glass Exetainer vials. Subsequent gas samples were collected after 7 h and 24 h. After each gas sampling, 12 mL of ambient air was injected into the chamber to maintain pressure equilibrium. N_2_O concentrations were quantified via gas chromatography (Agilent 7890A; Agilent Technologies, Palo Alto, CA, USA). The amount of N_2_O-N produced is given in ng N_2_O-N g^-1^ dw soil and were calculated as follows:

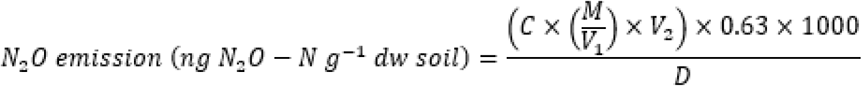

where C is the measured N_2_O concentration (µmol mol^-1^), M denotes the molar mass of N_2_O (44.01 µg µmol^-1^), V_1_ is the volume occupied by 1 mol of gas under standard conditions (24.45 L mol^-1^), V_2_ is the headspace volume (L), 0.63 represents the mass fraction of nitrogen in N_2_O, 1000 is a conversion factor and D is the soil dry weight enclosed inside the chamber. The percentage of N losses as N_2_O was calculated by averaging the maximum N_2_O produced in each treatment after each irrigation event and expressing this value as a fraction of the total N applied (117 µg N g^-1^ dw soil). *^15^N uptake in plant biomass*

To evaluate whether the addition of NIs improved plant nitrogen uptake, a ^15^N solution of (NH₄)₂SO₄ was applied five days before the final harvest. The ^15^N solution was injected into the planted and bare soil compartments at depths of 4 and 9 cm (3 and 2 mL from a 0.6 M solution were applied into the planted and bare soil compartments, respectively). At the final harvest, plant material was separated into roots and shoots. The roots were first scanned and images analyzed using RhizoVision (Seethepalli et al., 2024). Root morphology was determined based on total root length, average root diameter, specific root length, root density, and branching frequency. After drying, the biomass was weighed, and the plant material was then ground. To determine N and carbon concentrations, as well as the ^15^N isotopic composition in plant material, barley roots and shoots, 1-2 mg of ground plant tissue were analyzed using a thermochemical elemental analyzer (TC/EA, Thermo Fisher) coupled to an isotope ratio mass spectrometer (Delta V Advantage, Thermo Fisher).

### Microbial carbon dynamics and community composition

The soil microbial community plays a key role in plant growth as well as in several soil biogeochemical processes (Zhao et al., 2014; Zhao et al., 2025). Consequently, interactions between NIs and soil microbial communities not only affect NI efficacy but may also lead to unintended impacts on plant-microbe interactions. To assess the effect of NI addition on the efficiency of microbial carbon allocation, microbial biomass carbon (MBC), microbial growth, and respiration were measured at the final harvest. Microbial biomass carbon was determined by the chloroform-fumigation extraction in which fumigated and non-fumigated soil samples were extracted with 0.1 M K_2_SO_4_ (2 g soil in 15 mL K_2_SO_4_) and analyzed for dissolved organic carbon (DOC) on a TOC/N Analyzer (TOC-V CPH E200V/TNM-122 V, Shimadzu, Austria). Microbial biomass carbon was calculated as the difference in DOC between the fumigated and non-fumigated samples, using a correction factor of 0.45 (Vance et al., 1987; Jenkinson et al., 2004).

Microbial growth (μg C h^-1^ g^-1^ dw soil) was estimated through the incorporation of ^18^O from labelled water into DNA (Simon et al., 2020). For this, 500 mg of fresh soil were weighed into 1 mL cryovials, which were then placed open inside 27 mL glass vials. After airtight sealing, the soil was amended with ^18^O-labeled water (25µl of 97at% labelled ^18^O water; Campro Scientific). A set of samples taken from each treatment were amended with the same volume of DNAse free water and were used to determine ^18^O natural abundance. After 24h of incubation, labeled and unlabeled samples were stored at-80°C. DNA of both sets of samples was extracted using the DNeasy PowerSoil Pro kit (Qiagen, Hilden, Germany) according to the manufacturer’s instructions. The ^18^O isotopic composition, and total oxygen content of the DNA, were determined after drying a 50 µL aliquot of DNA extract for 24h at 60°C, followed by measurement on a Thermochemical Elemental Analyser (TC/EA Thermo Fisher) coupled via a Conflo III open split system (Thermo Fisher) to an Isotope Ratio Mass Spectrometer (IRMS, Delta V Advantage, Thermo Fisher).

Microbial respiration (μg C h^-1^ g^-1^ dw soil) was determined through concurrent measurements of CO_2_ during a 24-incubation period. For this, 5 mL gas samples were taken after ^18^O water amendment and at the end of the incubation period. The gas samples were then measured using an infrared gas analyzer (EGM-4 Environmental gas analyzer for CO_2_, PP Systems; Hertfordshire, UK). Microbial CUE was calculated as the ratio of the microbial growth rate to the sum of microbial growth and respiration rates.

To evaluate the effect of NI addition on the microbial community composition, bacterial and archaeal 16S rRNA gene V4 region gene amplicon sequencing was performed on the DNA extracted from the ^18^O water incubation. The 16S rRNA gene amplification and sequencing was carried out using the 515F/806R primers (Apprill et al., 2015; Parada et al., 2016) and on the Illumina MiSeq platform, at the Joint Microbiome Facility of the Medical University of Vienna and the University of Vienna JMF (project ID: JMF-2409-06). Amplicon sequence variants (ASVs) were inferred using the DADA2 pipeline (Callahan et al., 2016) following the recommended workflow. Taxonomy was assigned using the SILVA database SSU Ref NR 99 release 138.1 (Quast et al., 2013).

### Soil N pools quantification

To assess the residual effects of NIs on soil nitrogen pool concentrations and composition at the end of the experiment, soil total N, inorganic N species (NO_2_^-^, NO_3_^-^, NH_4_^+^), dissolved inorganic N (DIN), and dissolved organic N (DON) were determined. Soil from the planted and bare soil compartments was extracted with 0.1 M K_2_SO_4_ (2 g soil in 15 mL K_2_SO_4_), and the total N content was measured using a TOC/TN analyzer (TOC-V CPH E200V/TNM-122V; Shimadzu, Austria). The acidic VCl_3_/Griess reaction was used to determine NO_2_^-^ and NO_3_^-^ concentrations in the K_2_SO_4_ extracts (Griess-Romijn van Eck, 1966; Miranda et al., 2001), and a modified photometric indophenol reaction method was used to determine NH_4_^+^ (Kandeler and Gerber, 1988; Mulvaney, 1996). DIN was calculated by summing the NO_2_^-^, NO_3_^-^, and NH_4_^+^ concentrations, and DON was calculated by subtracting DIN from the total N content.

## Statistical analysis

Effects of treatment and compartment on the percentage of N loss (leachates and gaseous emissions) were assessed using a two-way ANOVA followed by Tukey’s HSD. Differences in root ^15^N content normalized to biomass, soil N pool concentrations, and microbial carbon-related processes among treatments were assessed using one-way ANOVA followed by Tukey’s HSD. Where assumptions of normality were not met, Welch’s ANOVA with Games–Howell post-hoc tests were applied (NO_2_^-^ concentrations and shoot ^15^N content normalized to biomass). Differences in plant biomass and root-to-shoot mass ratio among treatments were tested using Kruskal–Wallis followed by Dunn’s post hoc test. To test the effects of treatment, compartment, and sampling day on cumulative N losses in leachates, as well as differences in atom percent excess among treatments and between plant tissues, we fitted gamma-log generalized linear models (GLMs; glm). Pairwise comparisons were conducted using estimated marginal means (EMMs) with Bonferroni correction (emmeans, v1.11.1) (Lenth, 2025). All statistical analyses were performed in R version 4.4.3 (R Core Team, 2025) using Rstudio version 2025.09.2 (Posit Team, 2025).

Analyses of the 16S rRNA gene amplicon sequencing data were performed using the *phyloseq* 1.50.0 (McMurdie and Holmes, 2013), *microbiome* 1.28.0 (Lahti and Shetty, 2012), and *DESeq2* 1.46.0 (Love et al., 2014) packages. Alpha-diversity analysis was performed on rarefied datasets. To assess the effects of the NIs and compartment on microbial community composition, principal component analysis (PCA) based on an Aitchison distance matrix, and a PERMANOVA were performed. Finally, effects of NI application on 16S rRNA gene-based microbial communities of the planted and bare-soil compartments were assessed through genus-level differential relative abundance analyses, after excluding genera with an average relative abundance < 2% in any treatment.

## Results

### Nitrogen losses from the rootbox system

To evaluate the effects of BNI addition on the magnitude of N fertilizer losses, and to compare it to the effect of the SNI DMPP, the concentrations of mineral N in the leachates and the maximum N_2_O produced within 24 h after irrigation were determined for both planted and bare soil with and without NI addition. Across all treatments, in both planted and bare soil compartments, N was primarily lost via NO_3_^-^ leaching (ranging from 3.9 ± 0.3% to 8.8 ± 1.9% of the N applied, Fig. 1, Fig. S2), while NH_4_^+^ and NO_2_^-^ losses were negligible (NH_4_^+^: 0.02-0.81% of the N applied; NO_2_^-^: 0.05-0.8% of the N applied, Figure S2). Compared to the control, MBOA and BNI mixture application resulted in significantly greater NO_2_^-^ accumulation but lower NO_3_^-^ leaching, at specific time points (Fig. S2). N losses via leaching did not significantly differ between planted and bare soil or among NI treatments (Fig. 1A, B), but were on average higher in the controls without NI, with 9.4 ± 1.9% of the applied N lost via leaching in planted soil, and 8.3 ± 1.9% in bare soil.

**Fig. 1.**
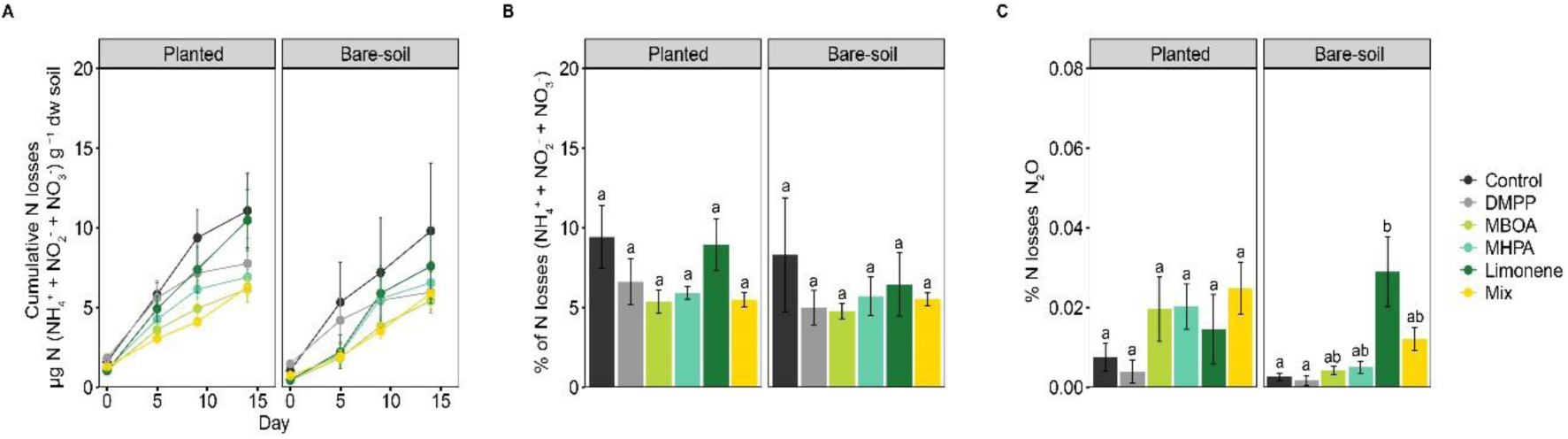
Mineral N losses via leaching and N_2_O emissions in planted and bare-soil compartments. **A** Cumulative N losses as NH_4_^+^, NO_2_^+^ and NO_3_^-^ in the leachates. Data are normalized per microgram N dry weight (dw) soil and presented as means ± SE (n = 4). **B** Fraction of applied N recovered in the leachates. The percentage of N losses in each treatment was calculated taking the cumulative losses at the final day of sampling (day14) and is presented as means ± SE (n =4) **C** Fraction of N applied emitted as N_2_O. The percentage of N lost as N_2_O was calculated by averaging the maximum N_2_O detected after each of the four irrigation events across the four replicates per treatment. Data are presented as means ± SE (n =8). Nitrogen fertilizer was applied on day 1 and day 10. Lowercase letters denote significant differences among treatments within each compartment. Results of post-hoc tests comparing treatments on each day within and between compartments are provided in Tables S1–S3.

In addition to losses via leaching, gaseous losses as N_2_O emissions were quantified at four timepoints. Regardless of compartment and NI type, N_2_O emissions represented the smallest N loss pathway (0.001–0.028% of the N applied, Fig. 1C), remaining lower than the already low NH_4_^+^ and NO_2_^-^ losses detected in the leachates. Interestingly, none of the NIs had a significant effect on N_2_O emissions from the planted soil. Yet, all BNI amendments led to a not statistically significant, but notable increase in N_2_O emission (Fig. 1C). In contrast, in bare soil limonene significantly increased N_2_O losses compared to the control. Overall, the three BNIs added and the mixture exhibited contrasting N_2_O dynamics (Fig. S3), with some showing sustained N_2_O accumulation (MBOA and MHPA) and others a rapid decline after peak concentrations (BNI mixture and limonene). These contrasting responses, indicate that the effect of BNIs on N_2_O production is compound-dependent, likely influenced by plant-soil interactions, and not trivial to constrain.

### Mineral N dynamics in the soil pore water

The mineral N concentration in the soil pore water reflects N directly available for plant uptake and microbial activity and provides deeper insight into the processes controlling mineral N dynamics in leachates. Similar as for the leachates, mineral N in the soil pore water was dominated by NO_3_^-^ (representing up to 93.5% and 89.4% from the N applied in the planted and bare soil, respectively. Fig. S4), while NO _2_^-^ represented between 0 - 8.5% of the N applied. NO_2_^-^ concentrations in the soil pore water of planted and bare soil were significantly higher after the application of MBOA and the BNI mixture, compared to the control (Fig. S4). Concurrently, NO_3_^-^ accumulation in the pore water was significantly reduced by MBOA and the BNI mixture in both the planted and bare soils. In contrast, DMPP significantly reduced NO_2_^-^ concentrations in both the planted and bare soil throughout the incubation, but had no effect on NO₃⁻ concentrations. Limonene only moderately changed pore water NO_2_^-^ and NO_3_^-^ concentrations, by significantly reducing them only on day 9 (Fig. S4). As already observed for the leachates, MHPA had no effect on NO_2_^-^ or NO_3_^-^ concentrations in pore water in either the planted or the bare soil compartments.

### Effect of NI application on plant biomass

The addition of nitrification inhibitors is expected to increase the availability of N for plant uptake, resulting in improved plant N nutrition and a potential increase in plant biomass (Liu et al., 2013; Abalos et al., 2014). Therefore, to determine whether NI application positively affected plant growth, shoot and root biomass were quantified after 14 days of plant growth. In contrast to our hypothesis, MBOA and MHPA additions did not affect barley shoot and root biomass compared to the controls without NI addition (Fig. 2A). DMPP induced a slight, although not statistically significant, increase in shoot biomass compared to the control (25.4% Fig 2A), with no effect on root biomass. Notably, the addition of limonene and the mixture of BNIs led to stunted growth, with significantly reduced shoot (65% reduction compared to the control in both treatments) and root biomass (60 and 45% reduction compared to the control in the limonene and mix treatments, respectively). Likewise, the limonene and the BNI mixture treatments led to an increase in the root-to-shoot ratio, indicating differences in biomass allocation (Fig. 2B). In contrast, MBOA, MHPA and DMPP had no statistically significant effect on the root-to-shoot ratio compared to the control. While there were no significant differences in the root tissue density among treatments (Fig. S5), the average root diameter, as well as the root branching frequency was significantly higher in the BNI mix treatment, explaining the greater root-to-shoot biomass ratio observed in this treatment. Finally, all treatments with the exception of DMPP resulted in a lower total N content in both shoot and root biomass, while MBOA and the BNI mix also lead to an increased C content in shoot biomass (Fig. S6).

**Fig. 2.**
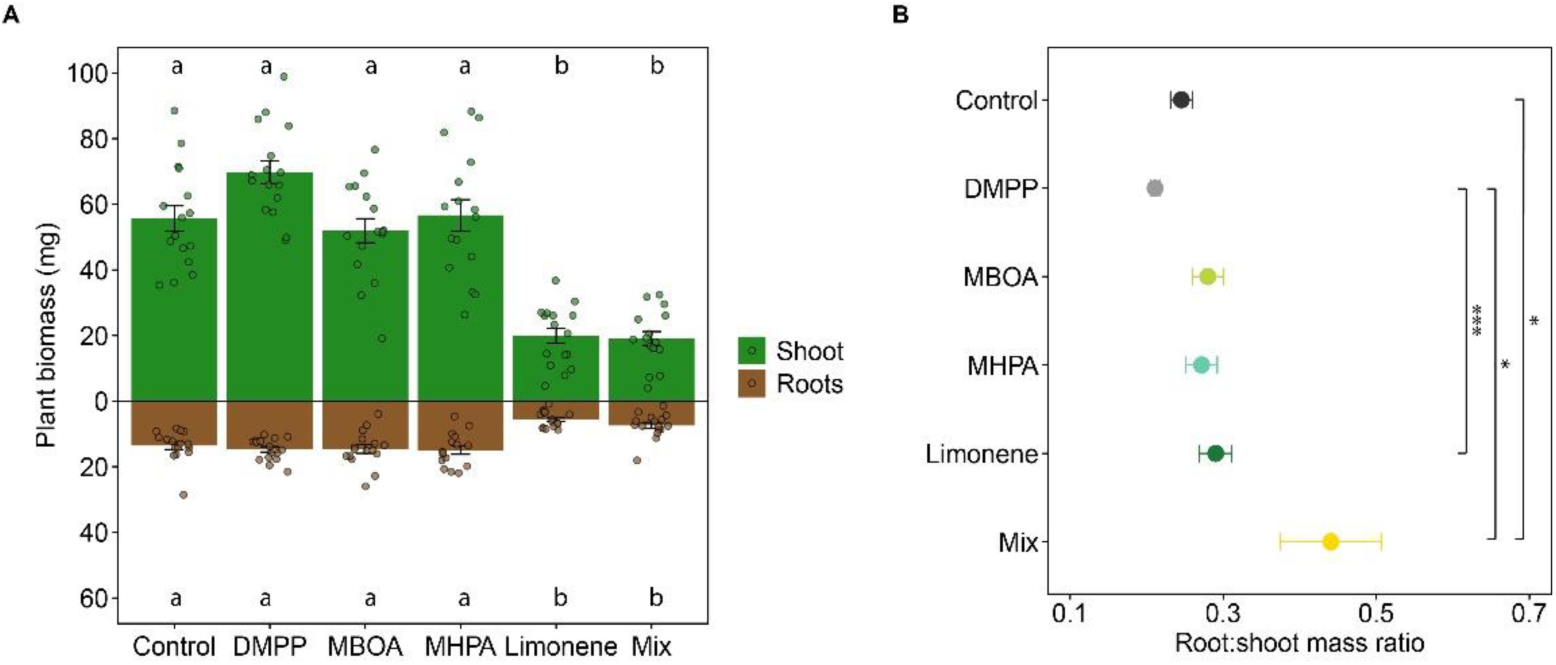
Effect of NIs on plant biomass and allocation. **A** Root and shoot dry weight and **B** root-to-shoot mass ratio for each NI treatment and the control. Data are presented as mean ± SE (n = 16; 4 plants × 4 rhizoboxes per treatment). Lowercase letters indicate significant differences among treatments. Results of post-hoc tests comparing treatment effect on plant biomass and root-to-shoot ratio are provided in Tables S4–S5.

### Effect of NI application on ^15^N allocation

We investigated the potential of BNIs to enhance fertilizer N uptake by tracing ^15^N labelled NH_4_^+^ from soil to plant shoot and roots. Six days after ^15^N addition, a higher ^15^N at% enrichment and total ^15^N content (mg ^15^N g^-1^ dw biomass) was observed in plant tissues compared to the ^15^N that remained in soil (Fig. 3. Fig. S7). Interestingly, DMPP and MBOA significantly reduced ¹⁵N enrichment (at%) in shoots compared to the roots, while the opposite was observed with limonene (Fig. 3A). Most of the NI treatments resulted in a similar ^15^N enrichment as the control, except for the BNI mixture, which significantly reduced ^15^N enrichment in shoots and roots (Figure. 3A). Despite plant-tissue and treatment-specific differences in ¹⁵N uptake, as well as an overall ¹⁵N enrichment similar to that of the untreated control, a significant reduction in ¹⁵N content per unit biomass was observed for all SNI and BNI treatments, with the exception of MPHA.

**Fig. 3.**
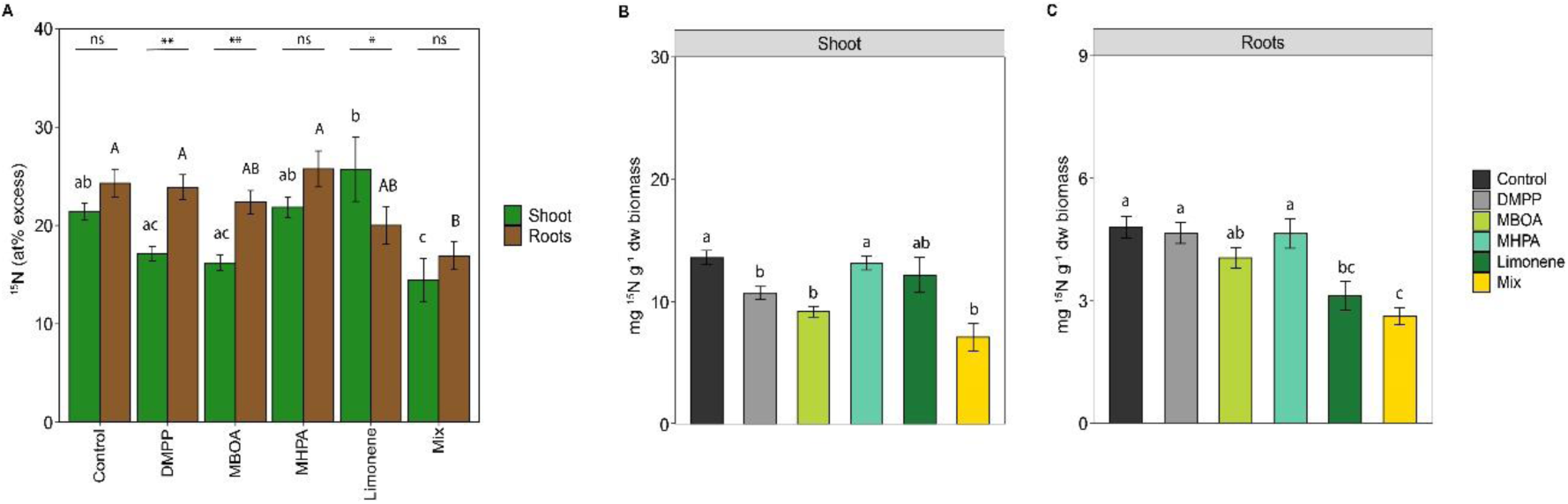
Plant ^15^N allocation. **A.** Atom percent excess in barley shoots and roots for NI treatment and the control. Significant differences in atom% excess among treatments are indicated by lowercase and uppercase letters for shoots and roots, respectively. Asterisks indicate significant differences between ^15^N in shoots and roots (* < 0.05, ** < 0.01). Total ^15^N content, normalized per gram of dry biomass, is shown for **B.** shoots and **C.** roots. Data are presented as means ± SE (n =16). Results of post-hoc tests comparing treatment effect on ^15^N allocation in the shoots and roots are provided in Tables S6–S7

### Effect of NI application on the soil N pools

In addition to assessing the effect of NI application on plant N uptake, their effect on overall soil N pools at the end of the experiment was evaluated. Specifically, soil total dissolved N (TDN), inorganic N species (NO_2_^-^, NO_3_^-^, NH_4_^+^), dissolved inorganic N (DIN), and dissolved organic N (DON) pools were quantified to evaluate whether changes in soil N availability and composition occurred. Total dissolved N ranged between 82 and 208 μg N g^-1^ dw soil, with DIN representing the largest fraction of the total N, while DON concentrations accounted for a maximum of 13.3% of TDN (Fig. 4). In the planted soil, both TDN and DIN concentrations were higher in the MHPA and BNI mixture treatments, while they remained comparable to the controls in DMPP, MBOA and limonene treatments (Fig. 4). Within the soil-bound DIN pool, NH_4_^+^ was the most abundant inorganic N form (between 23-115 μg N g^-1^ dw soil), followed by NO_3_^-^ (33.3-77 μg N g^-1^ dw soil), while NO_2_^-^ showed the lowest concentrations (0.06-33.5μg N g^-1^ dw soil). Notably, the presence or absence of plants did not have a significant effect on soil N pools.

**Fig. 4.**
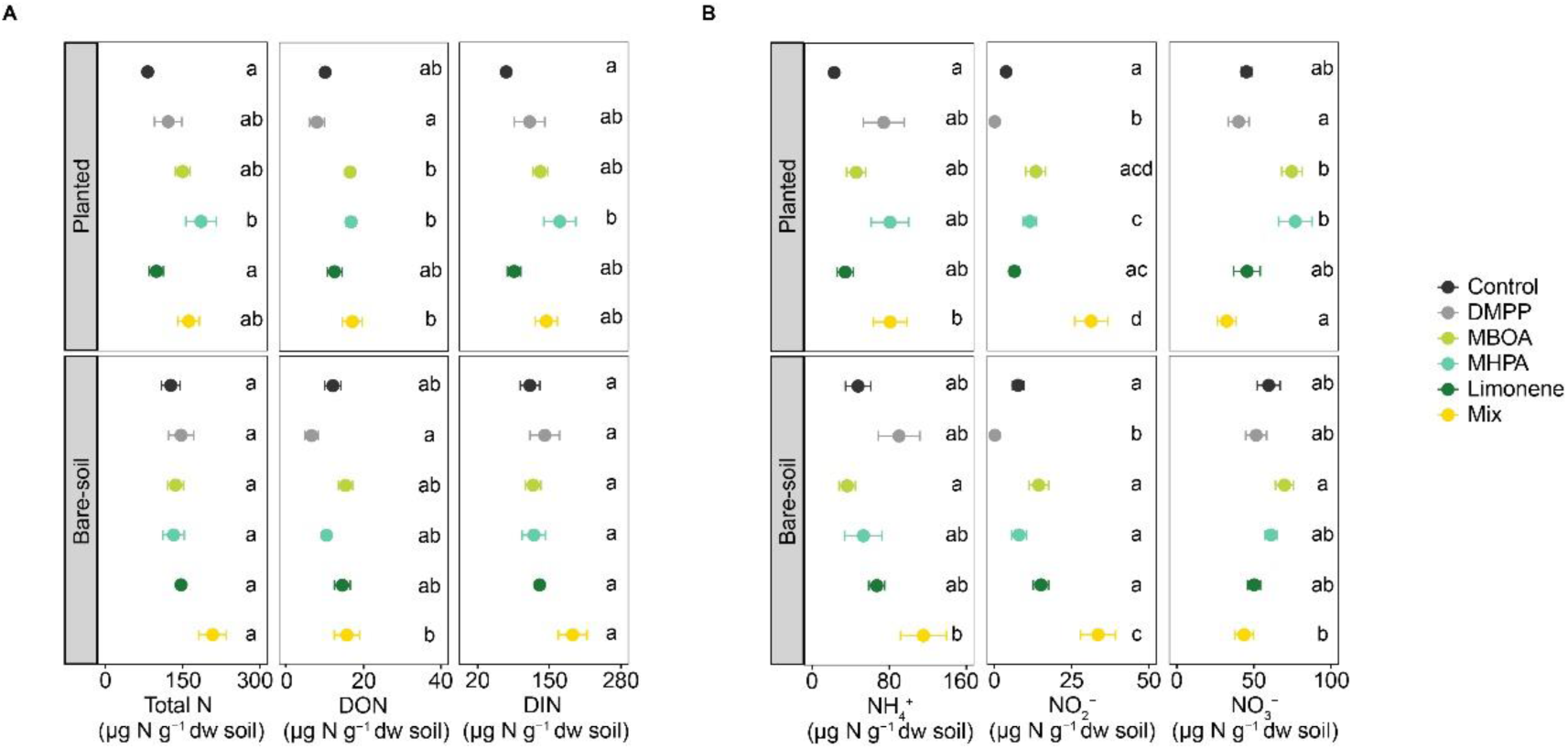
Effects of NIs on soil nitrogen pool concentrations and composition, measured 15 days after NI application. Concentrations of total dissolved N (TDN), inorganic N species (NO_2_^-^, NO_3_^-^, NH_4_^+^), dissolved inorganic N (DIN), and dissolved organic N (DON) are shown for each treatment in the planted and bare-soil compartments. Lowercase letters indicate significant differences among treatments within each compartment. Data are normalized per µg N dry weight (dw) soil and presented as means ± SE (n = 8). Results of post-hoc tests comparing treatment effects of each N pool within the planted and bare-soil compartments are provided in Table S8.

The NI additions had a significant effect on the concentrations of the different N forms. DMPP significantly increased soil NH_4_^+^ and reduced soil NO_2_^-^ concentrations compared to the control. DMPP application also resulted in a significant reduction of the DON concentration, while no significant change in soil NO_3_^-^ concentrations was observed. In contrast, MBOA application led to a significant accumulation of NO _2_^-^ in both planted and bare soils, as also observed in the leachates and pore water samples. Soil NO_3_^-^ concentrations were also significantly higher in this treatment, while soil NH_4_^+^ concentrations were not significantly different compared to the control. In the MHPA treatment, a significantly higher soil NO_3_^-^ concentration was observed, while for limonene treatments no significant effects on soil N pool concentrations were observed. Finally, the BNI mix application resulted in a significant increase in DIN, primarily due to a significant increase in NH_4_^+^ and NO_2_^-^ concentrations, which together ultimately increased the soil total N pool concentrations (Fig. 4). Overall, these results demonstrate that NI applications can substantially modify soil N pools, highlighting that NIs differ markedly in their modes of action, efficacy and off-target effects, which has important consequences for fertilizer N retention and overall soil N availability.

### Effect of NI application on soil microbial communities

As the soil microbial community plays a key role in plant growth, soil health and soil N cycling, the effects of NI additions on microbial biomass, growth and respiration were tested, and their effect on microbial community composition was determined. The presence of plants or BNI addition had no significant effect on carbon-related microbial processes, whereas the addition of the SNI DMPP significantly reduced microbial respiration, and since microbial growth remained unaffected, this reduction resulted in a significant increase in CUE in DMPP treated planted and bare soils (Fig. 5A)

**Fig. 5.**
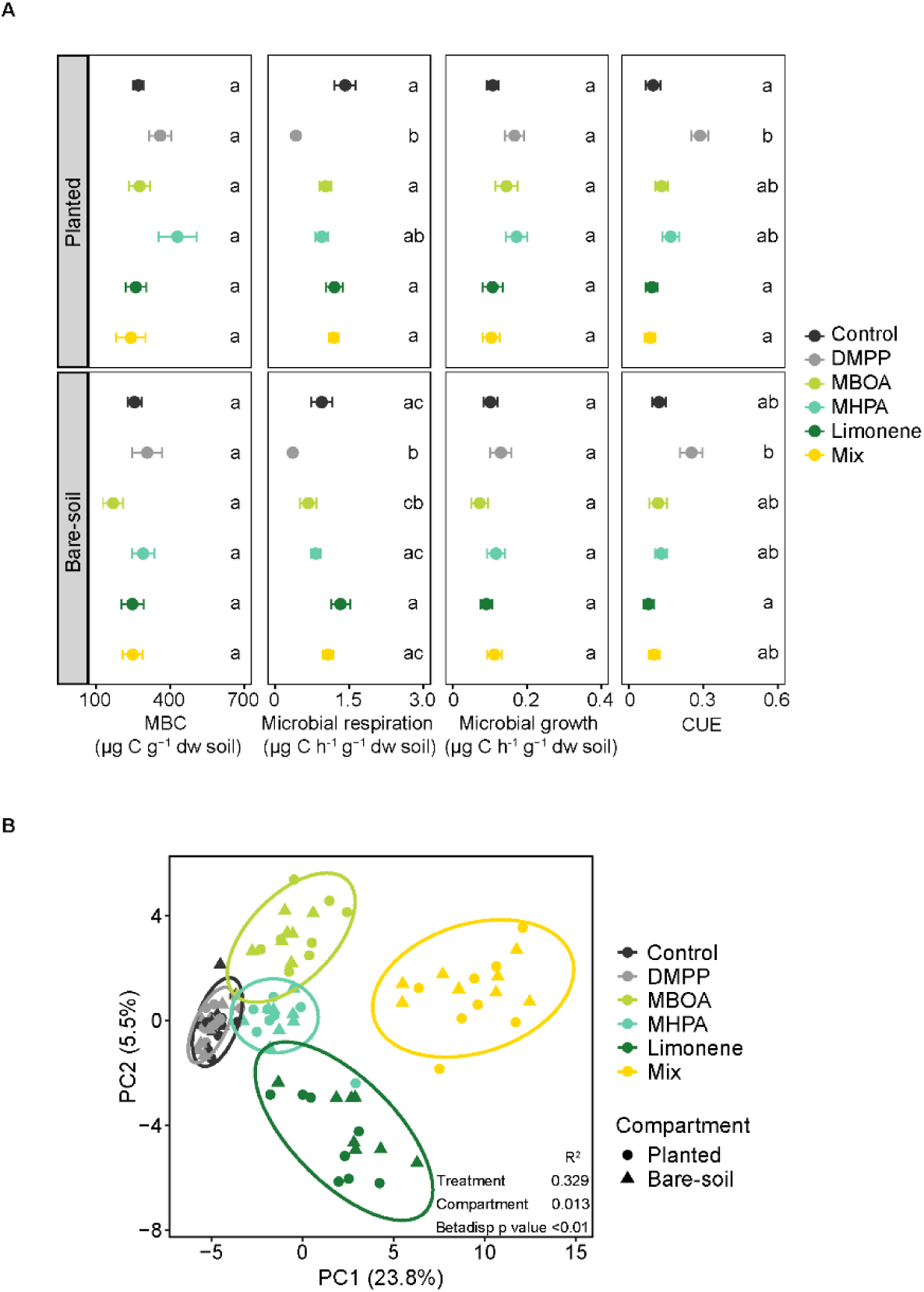
Microbial carbon processing and community composition responses to nitrification inhibitors. **A.** Effect of NIs addition on microbial biomass carbon (MBC), microbial respiration, growth, and carbon use efficiency (CUE) for the planted and bare-soil compartments. Lowercase letters indicate significant differences among treatments within each compartment. Data are normalized per µg N dry weight (dw) soil and presented as means ± SE (n = 8). **B.** Effect of NIs on the β-diversity of the bacterial community in the planted and bare-soil compartments. Principal component analysis at the genus-level based on an Aitchison distance matrix for the 16S rRNA gene-based communities. The variation explained by treatment and compartments, as well as the results from the multivariate analysis of dispersion (betadisper), are shown. Results of post-hoc tests comparing treatment effects of each microbial carbon-related process within the planted and bare-soil compartments are provided in Table S9.

While minor changes in carbon-related microbial processes were observed, BNI applications did substantially change soil microbial community composition and diversity (Fig 5B, Fig. S8 and S9). Notably, the presence of plants did not affect microbial community composition or diversity, and neither did the application of the SNI DMPP (Fig 5B). Community clustering according to BNI treatment indicates that each BNI selected for a distinct set of microbes or shifted the relative abundances of existing taxa in a compound specific manner (Fig 5B). In accordance, increases in the relative abundance of genera such as *Rhodococcus, Pseudomonas,* and *Lysobacter* were observed to be BNI specific, while an increased abundance of the genus *Novohierbaspirillum* was specific to the planted compartment (Fig. S9). Notably, the BNI mixture caused a substantial increase in the relative abundance of members of the *Rhodococcus* and *Pseudomonas* genera (∼21% and ∼15% increase in relative abundance, respectively Fig. S9). A similar increase in the relative abundance of members of the *Rhodococcus* genus was also observed after limonene application (∼30% increase in relative abundance Fig. S9). These compositional changes resulted in a significant reduction in microbial diversity in both the planted and bare soil (Fig. S8).

## Discussion

Nitrification is a major process through which fertilizer N is lost, which is why nitrification inhibitors are used to improve soil N retention for plant growth (Subbarao et al., 2012, 2013; Nardi et al., 2020; Buss et al., 2026). While BNIs have received a lot of attention in recent years and are widely discussed as an alternative to synthetic inhibitors, research has not yet fully integrated the effects of BNI addition in complex experimental systems involving interactions among plants, soil, and microbes (Kuppe and Postma, 2024; Buss et al., 2026). Using a rhizobox system with planted and unplanted alkaline soil, this study evaluated the effects of a widely used SNI (DMPP), three previously described BNIs, and a BNI mixture on plant N uptake, N losses, soil N pools, and the soil microbial community. Regardless of the NI treatment, mineral N losses in the rhizobox system occurred primarily as NO_3_^-^ leaching, which was also the predominant N form in the soil pore water (Fig. 1 and Fig. S2). Consistent with other cropping systems, NO_3_^−^ leaching represents a major pathway of N loss in barley cultivation (Richards et al., 1995; Vogeler et al., 2023). Considering that the soil mineral N concentrations were very low before N addition (0.39 ± 0.1 µg NH_4_^+^ g^-1^ dry weight (dw) soil and 15.19 ± 0.8 µg NO_3_^-^g^-1^ soil), NO_3_^-^ production appears to be the result of the oxidation of the applied fertilizer NH_4_^+^. Whether with barley or without, N losses in the leachates represented between 8 and 9.5% of the N applied in the untreated controls. SNI and BNI application in the rhizobox system reduced them to an average of 5.9%. It is worth noting that the leaching losses recorded here correspond only to the early growth period. However, if integrated over the entire growing season, cumulative N losses through leaching, as well as the difference between losses from the control and NI-treated soil, may have been higher.

Notably, in the planted soil all NI treatments, except for limonene, resulted in an increased soil total N content at the end of the experiment, with MHPA resulting in the highest increase (Fig. 4). The significant increase in total N following MHPA application was driven by a significant increase in both DON and DIN. Although this proposed BNI did not seem to effectively inhibit nitrification, and the inorganic N pool was therefore enriched mainly in NO_3_^-^at the end of the experiment soil NH_4_^+^ still accounted for ∼70% of the N applied. The concurrently increased NO_3_^-^ pool together with the increase in DON suggests that MHPA stimulated microbial N cycling, and a fraction of the soil NH_4_^+^ measured originated from the mineralization of organic matter rather than from the applied fertilizer NH_4_^+^. Similarly, increased gross mineralization and inorganic N immobilization have previously been observed in high-BNI genotypes of the tropical grass from which MHPA was originally isolated (Vázquez et al., 2020). The addition of a similar phenylpropanoid-type BNI together with organic amendments also increased gross mineralization in another alkaline soil, while having only minor effects on nitrification when applied without organic amendment (He et al., 2023).

A stimulating effect on DON in the planted soil was also observed following application of MBOA and the BNI mixture. However, in these treatments at the end of the experiment, NH_4_^+^ accounted for approximately 40% and 70% of the applied N, respectively, thus resulting in less efficient soil N retention than observed for MHPA. Application of the SNI DMPP did most effectively suppress nitrification based on soil DIN dynamics, but also resulted in a significant reduction in DON. This decrease in organic N, together with a significant reduction in microbial respiration, suggests that DMPP exerted an overall suppressive effect on soil microbial activity (Florio et al., 2016). Additionally, the final NH_4_^+^ pool was smaller (∼60% of the N applied) than in some BNI treatments. Taken together, these results demonstrate that SNI and BNI exert distinct effects on soil microbial communities and N transformations, and that understanding of these complex interactions is crucial for optimising N use efficiency in agricultural systems.

Several observations from soil microcosm incubations and pure-culture studies have demonstrated that DMPP effectively inhibits nitrification (Shi et al., 2016; Papadopoulou et al., 2020; Zhou et al., 2020; Bachtsevani et al., 2021; Bozal-Leorri et al., 2022), an effect we could also reproduce in this study. Still, DMPP treatment did not significantly reduce N losses, nor did it largely increase plant N uptake in our rhizobox system. While a significant reduction in NO_2_^-^ concentration compared to the untreated control was observed in the pore water of the DMPP treatment after the first irrigation, suggesting inhibition of nitrification, this effect diminished in later timepoints. It is likely that the large fractions of the water-soluble DMPP were leached out by irrigation from the root box system. Previously, higher DMPP mobility following rainfall, as well as reduced sorption in soils with dissolved organic carbon similar to that of the soil used here, have been observed (Marsden et al., 2016). In consequence, the imposed irrigation regime likely reduced DMPP persistence and, consequently, its efficacy in this alkaline soil. Such losses of DMPP to groundwater, or at least to deeper soil layers, are also likely to occur under field conditions, rendering its application ineffective.

Interestingly, application of MBOA, the BNI mixture and in some cases limonene led to significant NO_2_^-^ accumulation both in the leachates and in the pore water, accompanied with a significant reduction in NO_3_^-^ accumulation (Fig. S2 and S4). A reduction in O_2_ concentration, which may result from the high respiration rates of heterotrophs stimulated by the carbon provided by these compounds, might have indirectly inhibited nitrite oxidizing bacteria (NOB). However, a direct inhibition of NOB by the BNI is also possible. Typically the efficacy of newly discovered BNIs is assessed by their potential to reduce ammonia oxidation (Subbarao et al., 2006; Otaka et al., 2021, 2023; Kaur-Bhambra et al., 2022), while the effects of BNIs on NOBs have not been a major focus of study. Among the few studies that have examined this aspect, some have found greater sensitivity of NOB strains than AOB or AOA strains to BNIs (Kolovou et al., 2023). Given that NO_2-_ accumulation can stimulate aerobic N_2_O production in soils (Giguere et al., 2017), inhibition of NOBs may represent an important unintended negative effect of BNI application that should not be overlooked.

Our findings suggest that SNI and BNI modulates microbial soil N cycling in a more complex manner than anticipated or previously described. For instance, MHPA did not have a significant effect on nitrification, despite its in vitro potential to inhibit this process (Gopalakrishnan et al., 2007). However, its effect on soil microbial N cycling resulted in higher N stabilisation, which positively affects plant NH_4_^+^ availability. The stimulation of microbial N immobilization and the slower mobilization of organic N support temporal niche differentiation between the soil microbial community and the plant, thereby promoting a better long-term stabilization of soil N (Kuzyakov and Xu, 2013). Additionally, the slight but noteworthy differences observed here between planted and bare soil after BNI application reflect complex interactions between BNI activity, plant N uptake, root exudation, and soil microbial processes. Although these results are based only on the early phase of plant growth, they indicate that exogenous BNI addition affects primarily N remineralisation and DON pools, promoting NH_4_^+^ availability via soil N stabilisation. As this mode of action can potentially have an enhancing effect on plant growth, the whole plant-soil-microbe system should be routinely considered when assessing BNI efficacy. In line with our observations, a growing body of evidence suggests that an increase in microbial N immobilisation caused by BNIs application or exudation, rather than direct inhibition of ammonia oxidation, might explain the significant reduction in NO_3_^-^ accumulation observed in BNI efficacy studies. In a soil microcosm study, application of two BNIs resulted in a significant reduction in NO_3_^-^without a corresponding increase in residual NH_4_^+^, which the authors suggested may be due to increase in microbial immobilisation of NO_3_^-^ and/or NH_4_^+^ (Ma et al., 2021). Similarly, the low soil NO_3_^-^ content of high-BNI genotypes of tropical grasses has been associated with higher inorganic microbial N immobilisation rates (Karwat et al., 2017; Vázquez et al., 2020).

Under moderate to high N fertilization rates (40-170 kg N ha^-1^) the proportion of N applied recovered in spring barley typically ranges from 50-66%, reflecting the low intrinsic N use efficiency of this crop (Beatty et al., 2010; Anbessa and Juskiw, 2012). Plant preferences for N forms vary with soil properties, type of crop and environmental conditions (Elrys et al., 2025; Mao et al., 2025). However, studies with barley suggest that in general it uses both NH_4_⁺ and NO_3_^-^ and exhibits flexible, use of both forms (Richter et al., 1975; Kronzucker et al., 1999). Unlike other cereals in the same family, barley does not possess strong intrinsic BNI capacity (Subbarao et al., 2012, 2013). If so-called BNI compounds of other plants in fact primarily act by increasing microbial N immobilisation rates rather than by reducing nitrification rates, addition of an immobilisation stimulator like MHPA might have a positive effect of barley NUE without directly targeting nitrification.

An increase in mineral N availability via fertilisation, together with the C input provided by the BNIs, creates favourable conditions for heterotrophic growth and respiration, which increases microbial N immobilisation. Yet, these conditions also increase the likeliness of other undesirable N transformations to occur. For instance, limonene addition to the bare soil significantly increased N_2_O emissions compared with the control (Fig. 1C). Although N_2_O emissions represented the smallest form of N loss from the rhizobox system, irrigation events and NO_2_^-^ accumulation may have stimulated N_2_O production through both denitrification and nitrification. Previously, an increase in short-term N_2_O emissions, and the proportion of N_2_O production from denitrification was observed in a microcosms study where the BNI methyl 3-(4-hydroxyphenyl) propionate (MHPP) was added to an alkaline soil (Lan et al., 2023). A similar pattern was also observed in a low-pH soil microcosm study, where two BNIs applied at concentrations similar to those used here significantly increased cumulative N_2_O emissions relative to the control (Ma et al., 2021). Previously, it has been shown that the alleviation of C limitation rather than changes in soil NO_3_^-^ content due to BNIs played a more important role on N_2_O production and consumption (Florio et al., 2021). Readily available organic C sources are known to favor complete denitrification, and exert control over the abundance of different types of N_2_O reducing communities in agricultural soils (Surey et al., 2020; Maheshwari et al., 2023). It is in that context surprising that higher N_2_O emissions were observed in BNI amended planted than bare soils.

Overall, BNIs did not significantly affect N losses via leaching or gaseous emissions but modified microbial N transformations in soil in a compound - and time-specific manner. In some cases, these dynamics undermined their efficacy, suggesting that future BNI studies should account for the carbon-to-nitrogen balance and consider effects on N pools beyond those immediately relevant to nitrification. This would ensure that the prevention of N leaching does not inadvertently increase other environmentally harmful nitrogenous emissions. Interestingly, NI application primarily affected biomass allocation rather than overall plant growth, suggesting that NI-specific changes in nitrogen availability influenced internal plant resource partitioning. An increase in root-to-shoot mass ratio indicates plants competing for soil resources, while a decrease suggests that plants are able to allocate more energy to shoot growth, than into resource acquisition (Lopez et al., 2023). Thus, a favorable outcome in the root-to-shoot ratio, as observed for DMPP, MHPA and MBOA, should be considered during BNI suitability evaluations. Similarly, most NI treatments did not increase overall ¹⁵N enrichment relative to the control, but had an effect on the distribution of acquired N and, interestingly, reduced ¹⁵N content per unit biomass in most cases. Limonene and the BNI mixture resulted in a significant reduction in plant biomass as well as reduced ¹⁵N enrichment in both shoots and roots. In both cases, limonene likely reduced nutrient availability by decreasing soil water retention and promoting hydrocarbon-degrading taxa. In the long term, this may affect C cycling by altering SOC stability via rhizosphere priming effects (Zhou et al., 2024). Additionally, our results show that BNI mixtures do not necessarily produce additive effects.

## Conclusion

Over the last years, there have been increasing efforts to identify biological nitrification inhibitors from various plants. However, economically relevant crops such as barley do not seem to have a strong intrinsic BNI capacity (Subbarao et al., 2012, 2013). Consequently, the effects of exogenous addition of the proposed BNIs MHPA, MBOA, and limonene, and their mixture on barley growth were investigated. By using a rootbox system in which N transformations and changes in the soil microbial activity and community composition were assessed in planted and non-planted soil with and without NI addition. Our results demonstrate that while both SNIs and some BNIs have the potential to reduce N losses through leachates (DMPP, MBOA and BNI mixture), others suggested BNIs, can be detrimental to plant growth (limonene and the BNI mixture). Interestingly, all NI treatments except for MHPA resulted in lower plant N uptake, even if total plant biomass yield was increased, as observed for DMPP. BNI application altered soil N transformations and plant N allocation in a compound-specific manner, and our results provide new insights indicating that MHPA, and likely most BNIs, modulate microbial communities in a complex way that has not been previously reported. In sum, the alternative mode of N availability modulation via stimulation of microbial processes and N mineralisation observed here after the application of some BNIs might convey a less direct, but long-lasting effect on fertilizer N availability compared to currently used SNIs. Altogether, these findings highlight a promising new direction in BNI research, where BNI application could potentially stabilize N cycling in agricultural soils without necessarily inhibiting nitrification, while still promoting plant growth.

## Acknowledgements

We thank the Austrian Agency for Health and Food Safety (AGES) for facilitating the sampling sites. We are grateful to Julia Ramesmayer and Jasmin Schwarz for their technical support with the sequencing of the samples, as well as Joana Seneca for the pre-processing of the sequencing datasets.

