## Supplementary figures and tables for "Potential of exogenous biological nitrification inhibitor addition to improve soil nitrogen availability for crop growth"

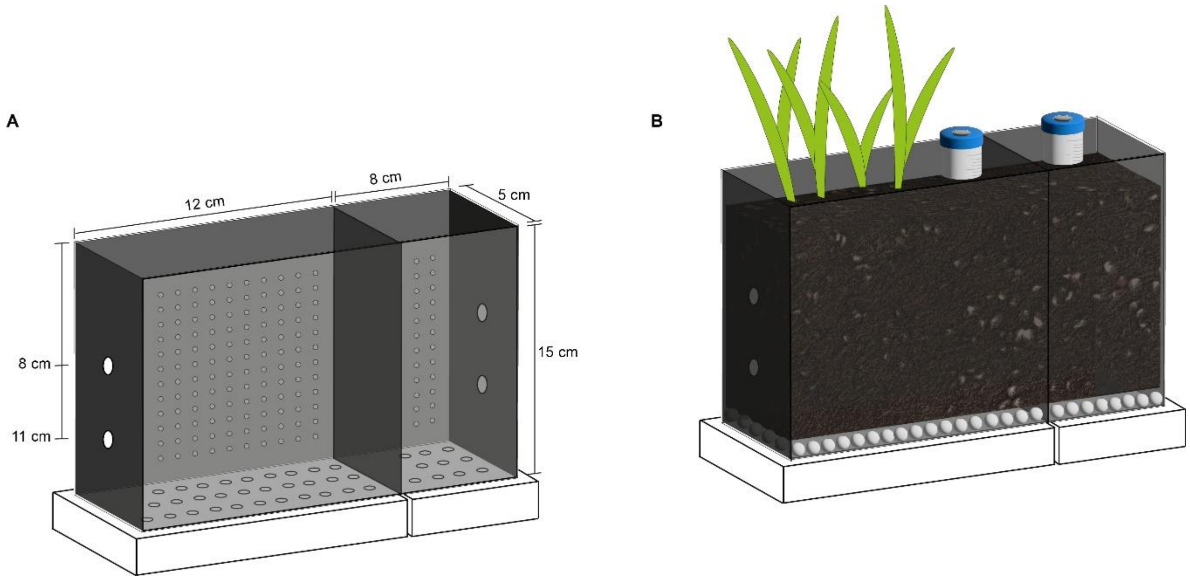


**Fig. S1.** Design of the rhizoboxes used in this study. **A** Empty rhizobox showing the dimensions of the planted and bare-soil compartment, which were separated by a solid strip isolating the compartments from each other. The depths at which soil pore water was sampled in both compartments are depicted. The bottom panels show the drainage holes (0.5-mm-diameter holes; spaced 1 cm apart) and the receiving tray for leachate collection. The back panel contained a 5 × 5 mm grid of holes (1 mm inner diameter), through which the $^{15}N$ solution, DMPP, and limonene were applied. **B** Schematic showing the arrangement of four plants per rhizobox in the planted compartment (4 rhizoboxes per treatment), as well as the gas chambers with screw caps (opened when not sampling) and the septum enabling continuous sampling during the first 24h post-irrigation. Glass beads used to improve drainage are also shown.

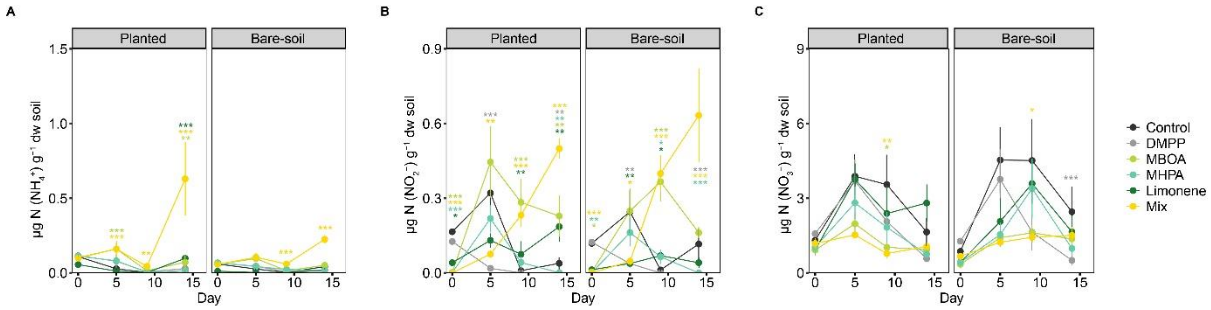


**Fig. S2.** Leachate mineral N concentration in the planted and bare-soil compartments. Effect of NI addition on the concentrations of **A.** $NH_4^+$, **B.** $NO_2^-$, and **C.** $NO_3^-$ detected in leachates from the planted and bare-soil compartments, expressed as μg N $g^{-1}$ dry soil. Data are presented as means ± SE (n = 4). Nitrogen fertilizer was applied on day 1 and day 10. Asterisks and the color denote significant differences of a specific treatment relative to the control on each day (* < 0.05, ** < 0.01, *** < 0.001). Post-hoc test results of differences on mineral N among treatments on each day for each compartment are provided in Tables S10-S12.

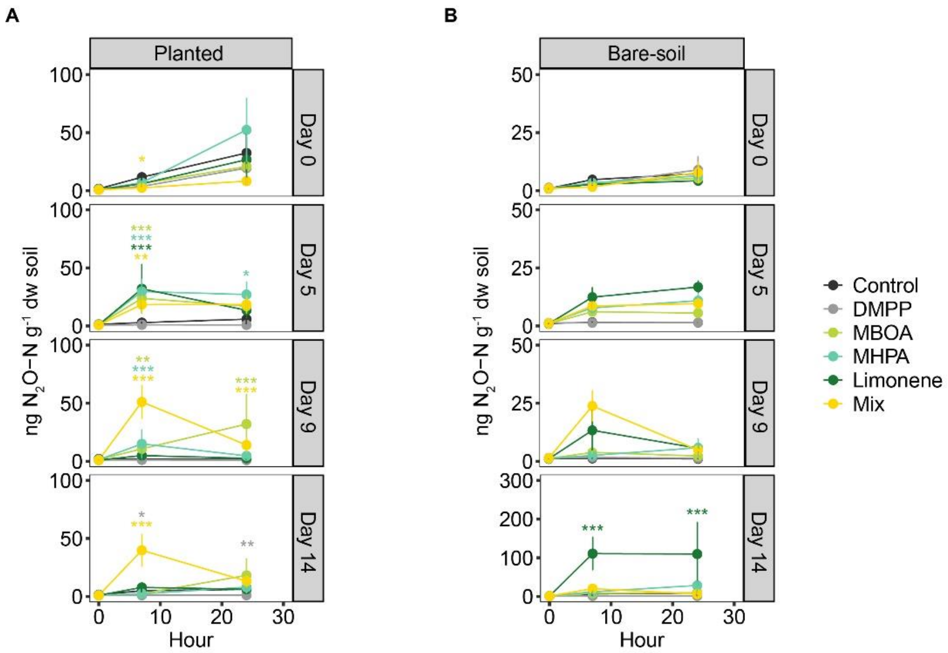


**Fig. S3.** Effect of NI addition on $N_2O$ emissions that occurred 24 h post-irrigation. $N_2O$ flux dynamics in **A.** planted and **B**. bare-soil compartments across treatments. Data are presented as means ± SE (n = 4). Nitrogen fertilizer was applied on day 1 and day 10. Asterisks and the color denote significant differences of a specific treatment relative to the control on each day (* < 0.05, ** < 0.01, *** < 0.001). Post-hoc test results of differences among treatments on each hour for each compartment are provided in Tables S13-S14.

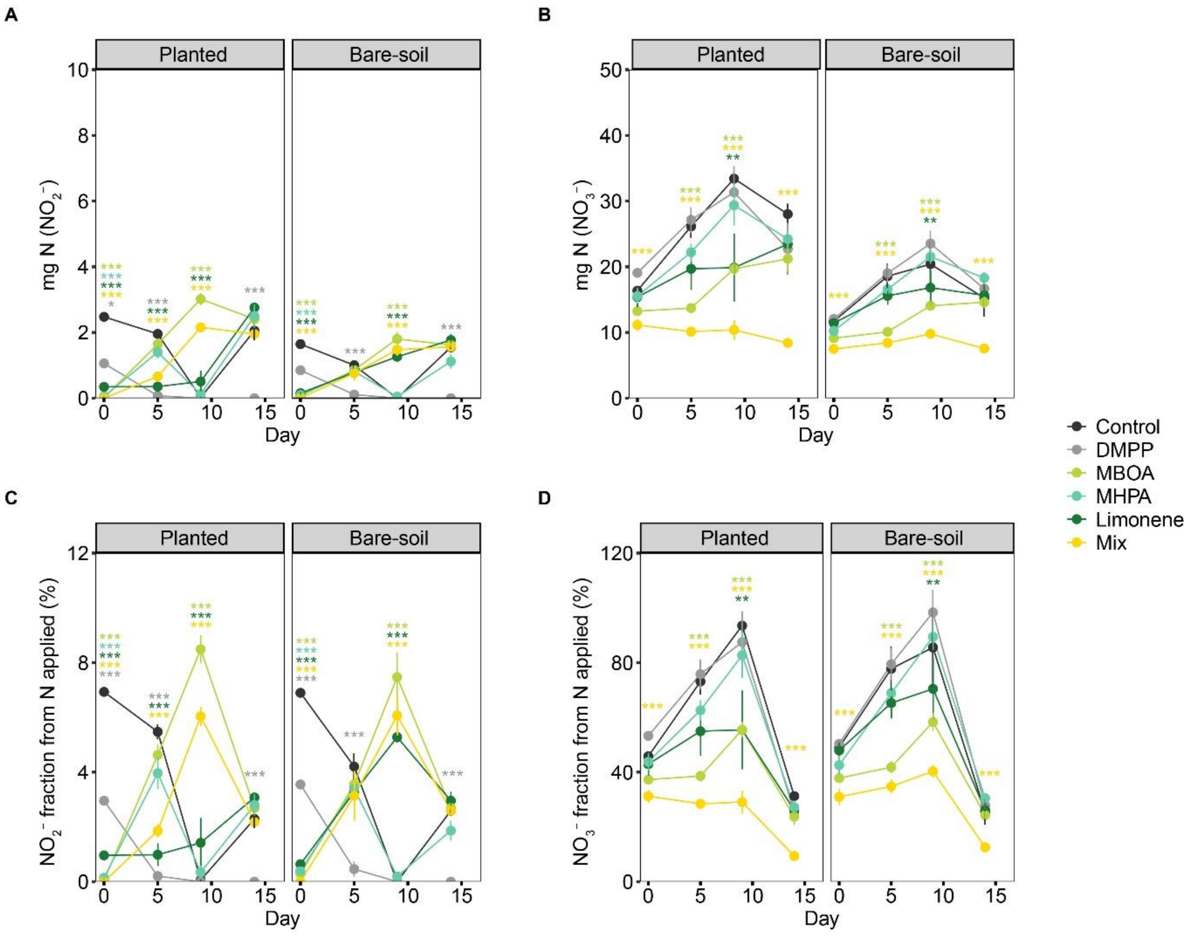


**Fig. S4.** Mineral N concentrations in the pore water of the planted and bare-soil compartments. Effect of NI application on the total **A.** $NO_2^-$ and **B.** $NO_3^-$ concentrations in the pore water of each compartment. Fractions of total N applied to each compartment detected as **C.** $NO_2^-$ and **D.** $NO_3^-$ in the pore water. Data are presented as means ± SE (n = 4). Nitrogen fertilizer was applied on day 1 and day 10. Asterisks and the color denote significant differences of a specific treatment relative to the control on each day (* < 0.05, ** < 0.01, *** < 0.001). Post-hoc test results of differences among treatments on each day for each compartment are provided in Tables S15-S18.

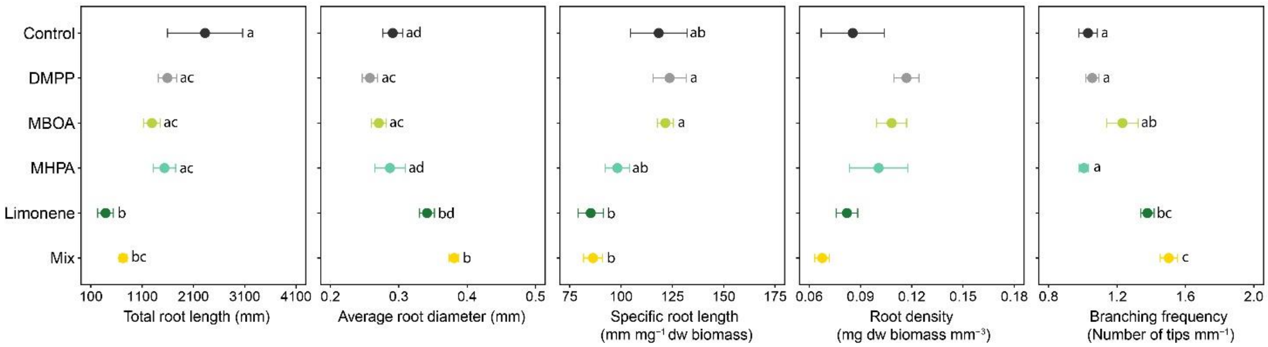


**Fig. S5.** Effect of NIs addition on root morphological traits. Total root length (mm), average root diameter (mm), specific root length (mm $mg^{-1}$ dw biomass), root density (mg dw biomass $mm^{-3}$), and branching frequency are shown for each NI treatment and the control. Data are presented as means ± SE (n = 8). Lowercase letters indicate significant differences among treatments.

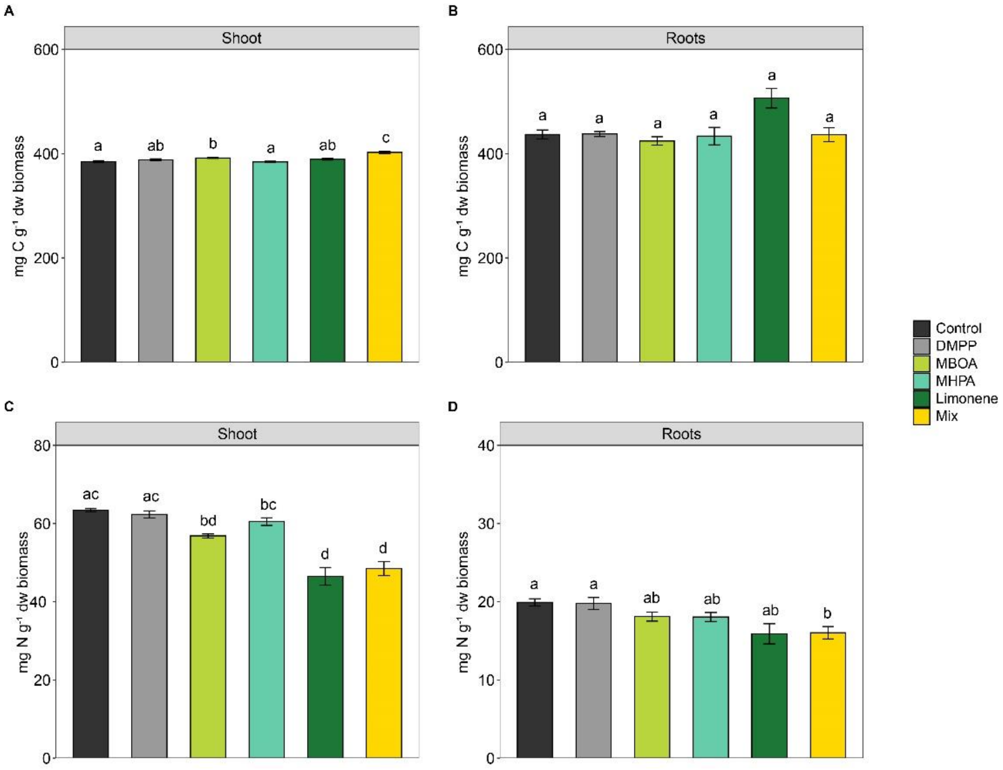


**Fig. S6.** Plant nitrogen and carbon content. Effect of NI addition on C content (mg C $g^{-1}$ dw biomass) in the **A.** shoots and **B.** roots. Effect of NI addition on N content (mg N $g^{-1}$ dw biomass) in the **C.** shoots and **D.** roots. Data are presented as means ± SE (n = 8). Lowercase letters indicate significant differences among treatments within each plant tissue.

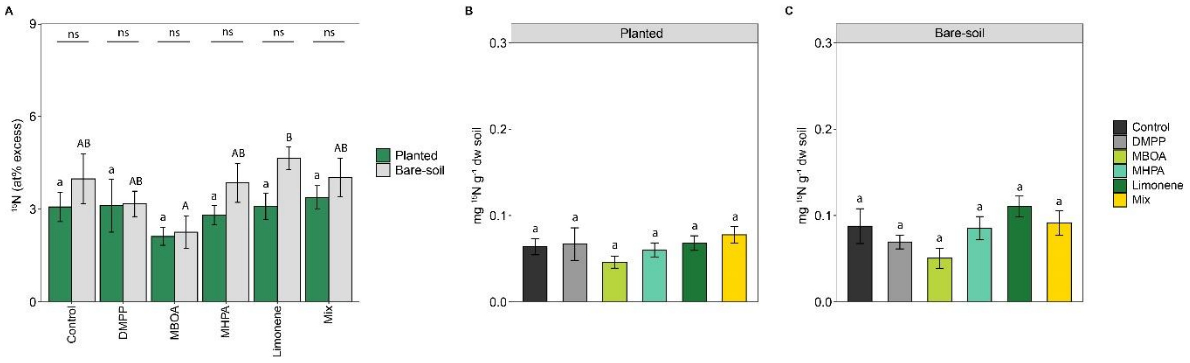


**Fig. S7.** Residual $^{15}$N content in the planted and bare-soil compartments. Effect of NI application on **A.** atom percent excess in the planted and bare-soil compartments. Significant differences in atom% excess among treatments are indicated by lowercase and uppercase letters for the planted and bare-soil compartments, respectively. Asterisks indicate significant differences in $^{15}$N content between compartments (ns: not significant). Total $^{15}$N content, normalized per gram of dry soil, is shown for **B.** planted and **C**. bare-soil compartments. Data is presented as means ± SE (n =8).

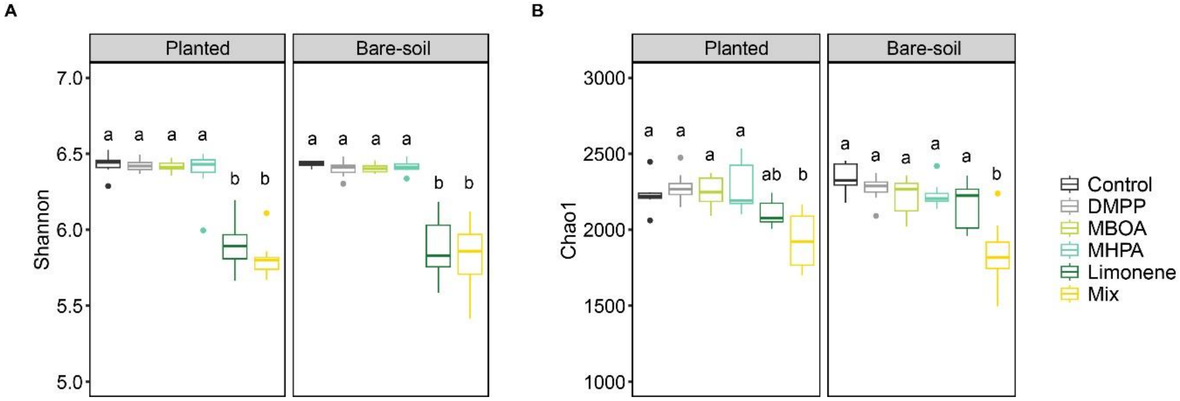


**Fig. S8.** Effect of NI addition on α-diversity indexes of the bacterial community in planted and bare soils. **A.** Shannon diversity index and **B.** Chao1 index for the 16S rRNA gene-based communities in soils from the planted and bare-soil compartments. In the boxplots, the median and the first and third quartiles are shown as the middle line and the lower and upper hinges, respectively. Whiskers extend to the smallest and largest values within 1.5 times the interquartile range, and outliers are shown as points. Lowercase letters indicate significant differences among treatments within each compartment.

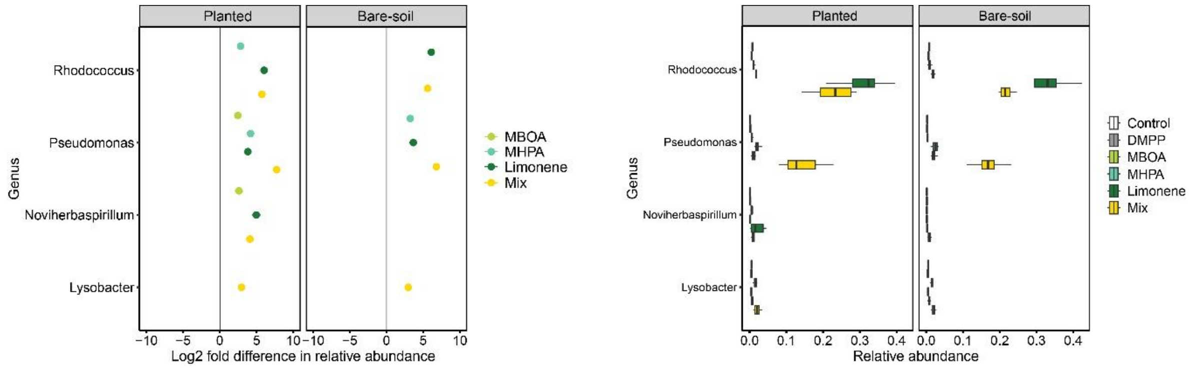


**Fig. S9.** Genus-level differential relative abundance changes for the 16S rRNA gene-based microbial community compositions from the planted and bare soils. Only genera with a relative abundance ≥ 0.02 in any of the treatments are depicted. In the boxplots, the median and the first and third quartiles are shown as the middle line and the lower and upper hinges, respectively. Whiskers extend to the smallest and largest values within 1.5 times the interquartile range, and outliers are shown as points. Data are presented as means ± SE (n = 8).

**Table S1:** Post-hoc test results for differences in cumulative N losses in leachates among treatments within the **(A)** planted and **(B)** bare-soil compartments, and **(C)** differences between compartments within each treatment. In all cases, pairwise comparisons of estimated marginal means (EMMs) from a gamma–log generalized linear model (GLM) were performed. Significant differences relative to the control and between treatments are shown in bold.

**A**

| **Comparison** | **Day 0** | | **Day 5** | | **Day 9** | | **Day 14** | |
|---|---|---|---|---|---|---|---|---|
| | **ratio** | **p adj** | **ratio** | **p adj** | **ratio** | **p adj** | **ratio** | **p adj** |
| Control-DMPP | 0.873 | 0.998 | 1.038 | 1.000 | 1.310 | 0.949 | 1.426 | 0.851 |
| Control- Limonene | 1.550 | 0.700 | 1.182 | 0.994 | 1.270 | 0.969 | 1.057 | 1.000 |
| Control- MBOA | 1.543 | 0.710 | 1.613 | 0.617 | 1.899 | 0.287 | 1.794 | 0.392 |
| Control- MHPA | 1.382 | 0.894 | 1.365 | 0.909 | 1.527 | 0.730 | 1.602 | 0.631 |
| Control- Mix | 1.233 | 0.983 | 1.908 | 0.280 | 2.287 | 0.077 | 1.760 | 0.430 |
| DMPP- Limonene | 1.777 | 0.411 | 1.139 | 0.998 | 0.970 | 1.000 | 0.741 | 0.921 |
| DMPP- MBOA | 1.768 | 0.421 | 1.554 | 0.695 | 1.450 | 0.824 | 1.258 | 0.974 |
| DMPP- MHPA | 1.583 | 0.656 | 1.315 | 0.945 | 1.166 | 0.996 | 1.124 | 0.999 |
| DMPP- Mix | 1.413 | 0.864 | 1.838 | 0.345 | 1.746 | 0.446 | 1.234 | 0.982 |
| Limonene- MBOA | 0.995 | 1.000 | 1.364 | 0.909 | 1.495 | 0.770 | 1.697 | 0.506 |
| Limonene- MHPA | 0.891 | 0.999 | 1.155 | 0.997 | 1.203 | 0.990 | 1.516 | 0.744 |
| Limonene- Mix | 0.795 | 0.975 | 1.614 | 0.615 | 1.801 | 0.384 | 1.665 | 0.547 |
| MBOA- MHPA | 0.895 | 0.999 | 0.846 | 0.994 | 0.804 | 0.980 | 0.893 | 0.999 |
| MBOA- Mix | 0.799 | 0.977 | 1.183 | 0.994 | 1.204 | 0.990 | 0.981 | 1.000 |
| MHPA -Mix | 0.893 | 0.999 | 1.398 | 0.879 | 1.497 | 0.768 | 1.098 | 1.000 |

**B**

| **Comparison** | **Day 0** | | **Day 5** | | **Day 9** | | **Day 14** | |
|---|---|---|---|---|---|---|---|---|
| | **ratio** | **p adj** | **ratio** | **p adj** | **ratio** | **p adj** | **ratio** | **p adj** |
| Control-DMPP | 0.713 | 0.994 | 1.265 | 0.988 | 1.322 | 0.941 | 1.642 | 0.578 |
| Control- Limonene | 2.290 | 0.748 | 2.394 | 0.183 | 1.222 | 0.986 | 1.292 | 0.959 |
| Control- MBOA | 2.516 | 0.498 | 2.880 | 0.057 | 1.871 | 0.312 | 1.809 | 0.375 |
| Control- MHPA | 2.115 | 0.626 | 2.563 | 0.123 | 1.295 | 0.957 | 1.496 | 0.770 |
| Control- Mix | 1.431 | 0.976 | 2.758 | 0.076 | 2.034 | 0.186 | 1.661 | 0.553 |
| DMPP- Limonene | 3.212 | 0.394 | 1.892 | 0.294 | 0.925 | 1.000 | 0.787 | 0.969 |
| DMPP- MBOA | 3.529 | 0.164 | 2.276 | 0.080 | 1.416 | 0.861 | 1.102 | 1.000 |
| DMPP- MHPA | 2.967 | 0.216 | 2.025 | 0.192 | 0.980 | 1.000 | 0.911 | 1.000 |
| DMPP- Mix | 2.007 | 0.696 | 2.180 | 0.113 | 1.539 | 0.715 | 1.012 | 1.000 |
| Limonene- MBOA | 1.099 | 1.000 | 1.203 | 0.990 | 1.531 | 0.725 | 1.400 | 0.877 |

| | | | | | | | | |
|---|---|---|---|---|---|---|---|---|
| Limonene- MHPA | 0.924 | 1.000 | 1.071 | 1.000 | 1.059 | 1.000 | 1.158 | 0.997 |
| Limonene- Mix | 0.625 | 0.924 | 1.152 | 0.997 | 1.664 | 0.549 | 1.286 | 0.962 |
| MBOA- MHPA | 0.841 | 0.997 | 0.890 | 0.999 | 0.692 | 0.830 | 0.827 | 0.989 |
| MBOA- Mix | 0.569 | 0.653 | 0.958 | 1.000 | 1.087 | 1.000 | 0.918 | 1.000 |
| MHPA -Mix | 0.676 | 0.791 | 1.076 | 1.000 | 1.571 | 0.673 | 1.110 | 0.999 |

**C**

| Comparison | Day 0 | | Day 5 | | Day 9 | | Day 14 | |
|---|---|---|---|---|---|---|---|---|
| | **ratio** | **p adj** | **ratio** | **p adj** | **ratio** | **p adj** | **ratio** | **p adj** |
| Control Bare- Planted | 0.660 | 0.389 | 0.916 | 0.814 | 0.767 | 0.385 | 0.886 | 0.690 |
| DMPP Bare- Planted | 0.808 | 0.658 | 0.751 | 0.348 | 0.760 | 0.368 | 0.769 | 0.389 |
| Limonene Bare- Plant | 0.447 | 0.096 | **0.452** | **0.010** | 0.797 | 0.456 | 0.725 | 0.291 |
| MBOA Bare- Planted | **0.405** | **0.016** | **0.513** | **0.030** | 0.779 | 0.411 | 0.878 | 0.670 |
| MHPA Bare- Planted | **0.431** | **0.006** | **0.488** | **0.020** | 0.905 | 0.744 | 0.949 | 0.863 |
| Mix Bare- Planted | 0.569 | 0.066 | 0.634 | 0.135 | 0.863 | 0.628 | 0.939 | 0.835 |

**Table S2:** Post-hoc test results for differences in the percentage of N losses in leachates among treatments within the planted and bare-soil compartments. Tukey HSD comparisons were done after a two-way ANOVA. Significant differences relative to the control and between treatments are shown in bold.

| Comparison | Planted | | Bare-soil | |
|---|---|---|---|---|
| | **diff** | **p adj** | **diff** | **p adj** |
| DMPP-Control | -0.365 | 0.984 | -0.333 | 0.992 |
| Limonene-Control | -0.043 | 1.000 | -0.137 | 1.000 |
| MBOA-Control | -0.541 | 0.806 | -0.313 | 0.995 |
| MHPA-Control | -0.417 | 0.958 | -0.200 | 1.000 |
| Mix-Control | -0.495 | 0.878 | -0.161 | 1.000 |
| Limonene-DMPP | 0.322 | 0.994 | 0.196 | 1.000 |
| MBOA-DMPP | -0.176 | 1.000 | 0.020 | 1.000 |
| MHPA-DMPP | -0.052 | 1.000 | 0.133 | 1.000 |
| Mix-DMPP | -0.130 | 1.000 | 0.172 | 1.000 |
| MBOA-Limonene | -0.498 | 0.875 | -0.176 | 1.000 |
| MHPA-Limonene | -0.374 | 0.981 | -0.063 | 1.000 |
| Mix-Limonene | -0.452 | 0.929 | -0.024 | 1.000 |
| MHPA-MBOA | 0.124 | 1.000 | 0.113 | 1.000 |
| Mix-MBOA | 0.045 | 1.000 | 0.152 | 1.000 |
| Mix-MHPA | -0.079 | 1.000 | 0.039 | 1.000 |

**Table S3:** Post-hoc test results comparing the percentage of N lost as $N_2O$ emissions among treatments within planted and bare-soil compartments. Tukey HSD comparisons were performed after a two-way ANOVA. Significant differences relative to the control and between treatments are shown in bold.

| Comparison | Planted | | Bare-soil | |
|---|---|---|---|---|
| | **Diff.** | **p adj** | **Diff.** | **p adj** |
| DMPP- Control | -0.004 | 1.000 | -0.001 | 1.000 |
| Limonene- Control | 0.007 | 0.998 | **0.026** | **0.042** |
| MBOA- Control | 0.012 | 0.893 | 0.002 | 1.000 |
| MHPA- Control | 0.013 | 0.853 | 0.002 | 1.000 |
| Mix- Control | 0.017 | 0.480 | 0.010 | 0.975 |
| Limonene - DMPP | 0.011 | 0.949 | **0.027** | **0.031** |
| MBOA- DMPP | 0.016 | 0.618 | 0.003 | 1.000 |
| MHPA- DMPP | 0.016 | 0.556 | 0.003 | 1.000 |
| Mix- DMPP | 0.021 | 0.214 | 0.010 | 0.953 |
| MBOA- Limonene | 0.005 | 1.000 | -0.025 | 0.071 |
| MHPA- Limonene | 0.006 | 1.000 | -0.024 | 0.092 |
| Mix- Limonene | 0.010 | 0.958 | -0.017 | 0.515 |
| MHPA- MBOA | 0.001 | 1.000 | 0.001 | 1.000 |
| Mix- MBOA | 0.005 | 1.000 | 0.008 | 0.994 |
| Mix- MHPA | 0.005 | 1.000 | 0.007 | 0.998 |

**Table S4:** Post-hoc test results comparing the effect of NI addition on shoot and root biomass. A Kruskal-Wallis test followed by Dunn's post hoc test were performed. Significant differences are shown in bold.

| Comparison | Shoot | | Root | |
|---|---|---|---|---|
| | **Statistic** | **p adj** | **Statistic** | **p adj** |
| Control-DMPP | 1.62 | 1 | 0.81 | 1 |
| Control- Limonene | **-4.32** | **< 0.001** | **-4.11** | **< 0.001** |
| Control- MBOA | -0.36 | 1 | 0.58 | 1 |
| Control- MHPA | 0.102 | 1 | 0.84 | 1 |
| Control- Mix | **-4.46** | **< 0.001** | **-3.16** | **0.0236** |
| DMPP- Limonene | **-5.95** | **< 0.001** | **-4.93** | **< 0.001** |
| DMPP- MBOA | -1.99 | 0.695 | -0.23 | 1 |
| DMPP- MHPA | -1.52 | 1 | 0.03 | 1 |
| DMPP- Mix | **-6.09** | **< 0.001** | **-3.98** | **0.001** |
| Limonene- MBOA | 3.95 | 0.001 | **4.70** | **< 0.001** |
| Limonene- MHPA | **4.42** | **< 0.001** | **4.96** | **< 0.001** |
| Limonene- Mix | -0.14 | 1 | 0.95 | 1 |
| MBOA- MHPA | 0.47 | 1 | 0.26 | 1 |
| MBOA- Mix | **-4.09** | **< 0.001** | **-3.74** | **0.003** |
| MHPA -Mix | **-4.56** | **< 0.001** | **-4.01** | **< 0.001** |

**Table S5:** Post-hoc test results comparing the effect of NI additon on shoot-to-root mass ratio. A Kruskal-Wallis test followed by Dunn's post hoc test were performed. Significant differences are shown in bold.

| Comparison | Statistic | p adj |
|---|---|---|
| Control-DMPP | -1.62 | 1 |
| Control- Limonene | 1.53 | 1 |
| Control- MBOA | 1.15 | 1 |
| Control- MHPA | 0.72 | 1 |
| Control- Mix | **3.06** | **0.033** |
| DMPP- Limonene | **3.15** | **0.024** |
| DMPP- MBOA | **2.77** | **0.083** |
| DMPP- MHPA | **2.35** | **0.283** |
| DMPP- Mix | **4.68** | **< 0.001** |
| Limonene- MBOA | -0.38 | 1 |
| Limonene- MHPA | -0.80 | 1 |
| Limonene- Mix | 1.53 | 1 |
| MBOA- MHPA | -0.42 | 1 |
| MBOA- Mix | 1.91 | 0.842 |
| MHPA -Mix | 2.34 | 0.293 |

**Table S6:** Post-hoc test results for differences in at% excess among treatments **A.** within each plant tissue and **B.** between plant tissues within each treatment. Pairwise comparisons of estimated marginal means (EMMs) from a gamma–log generalized linear model (GLM) were performed. Significant differences relative to the control and between treatments are shown in bold.

**A.**

| Comparison | Shoot | | Root | |
|---|---|---|---|---|
| | **ratio** | **p adj** | **ratio** | **p adj** |
| Control-DMPP | 1.250 | 0.352 | 1.017 | 1.000 |
| Control- Limonene | 0.832 | 0.611 | 1.213 | 0.601 |
| Control- MBOA | 1.322 | 0.133 | 1.086 | 0.979 |
| Control- MHPA | 0.980 | 1.000 | 0.943 | 0.995 |
| Control- Mix | **1.481** | **0.009** | **1.435** | **0.026** |
| DMPP- Limonene | **0.666** | **0.008** | 1.193 | 0.692 |
| DMPP- MBOA | 1.057 | 0.996 | 1.068 | 0.992 |
| DMPP- MHPA | 0.784 | 0.256 | 0.927 | 0.984 |
| DMPP- Mix | 1.185 | 0.673 | **1.411** | **0.040** |
| Limonene- MBOA | **1.588** | **0.001** | 0.895 | 0.945 |
| Limonene- MHPA | 1.177 | 0.722 | 0.777 | 0.301 |
| Limonene- Mix | **1.779** | **0.000** | 1.183 | 0.759 |
| MBOA- MHPA | 0.741 | 0.087 | 0.868 | 0.816 |
| MBOA- Mix | 1.120 | 0.918 | 1.321 | 0.175 |
| MHPA -Mix | **1.511** | **0.005** | **1.521** | **0.005** |

**B.**

| Comparison | ratio | p adj |
|---|---|---|
| Control Root-Shoot | 1.134 | 0.262 |
| DMPP Root-Shoot | **1.394** | **0.003** |
| Limonene Root-Shoot | **0.778** | **0.046** |
| MBOA Root-Shoot | **1.380** | **0.005** |
| MHPA Root-Shoot | 1.179 | 0.144 |
| Mix Root-Shoot | 1.171 | 0.182 |

**Table S7:** Post-hoc test results for differences in $^{15}N$ content (mg $^{15}N$ $g^{-1}$ dw biomass) among treatments in **A.** shoots and **B.** roots. Welch's ANOVA followed by the Games-Howell post hoc test was performed to test for significant differences in shoot $^{15}N$ content. Tukey HSD comparisons were performed following a one-way ANOVA to test for significant differences root $^{15}N$ content. Significant differences relative to the control and between treatments are shown in bold.

**A.**

| Comparison | Estimate | p adj |
|---|---|---|
| Control-DMPP | **-2.87** | **0.011** |
| Control- Limonene | -1.40 | 0.952 |
| Control- MBOA | **-4.39** | **<0.001** |
| Control- MHPA | -0.43 | 0.993 |
| Control- Mix | **-6.48** | **<0.001** |
| DMPP- Limonene | 1.46 | 0.940 |
| DMPP- MBOA | -1.52 | 0.248 |
| DMPP- MHPA | **2.43** | **0.033** |
| DMPP- Mix | -3.61 | 0.089 |
| Limonene- MBOA | -2.99 | 0.458 |
| Limonene- MHPA | 0.96 | 0.990 |
| Limonene- Mix | -5.07 | 0.125 |
| MBOA- MHPA | **3.95** | **<0.001** |
| MBOA- Mix | -2.09 | 0.552 |
| MHPA -Mix | **-6.04** | **0.002** |

**B.**

| Comparison | Diff. | p adj |
|---|---|---|
| Control-DMPP | -0.138 | 0.999 |
| Control- Limonene | **-1.681** | **0.003** |
| Control- MBOA | -0.753 | 0.452 |
| Control- MHPA | -0.158 | 0.998 |
| Control- Mix | **-2.186** | **<0.001** |
| DMPP- Limonene | **-1.543** | **0.008** |
| DMPP- MBOA | -0.614 | 0.670 |
| DMPP- MHPA | -0.020 | 1.000 |
| DMPP- Mix | **-2.047** | **<0.001** |
| Limonene- MBOA | 0.928 | 0.301 |
| Limonene- MHPA | **1.522** | **0.010** |

| | | |
|---|---|---|
| Limonene- Mix | -0.504 | 0.872 |
| MBOA- MHPA | 0.594 | 0.700 |
| MBOA- Mix | **-1.432** | **0.014** |
| MHPA -Mix | **-2.027** | **<0.001** |

**Table S8:** Post-hoc test results for differences in N pools concentrations (Total N, $NO_2^-$, $NO_3^-$, $NH_4^+$, dissolved inorganic N (DIN), and dissolved organic N (DON)) in the **A.** planted and **B.** bare-soil compartment. Tukey HSD comparisons were performed following a one-way ANOVA to test for significant differences in total N, $NO_3^-$, $NH_4^+$, DIN, and DON within each compartment. Welch's ANOVA followed by the Games-Howell post hoc test was performed to test for significant differences in $NO_2^-$. Significant differences relative to the control and between treatments are shown in bold.

**A.**

| Comparison | Total N | | NO2- | | NO3-- | | NH4+ | | DIN | | DON | |
|---|---|---|---|---|---|---|---|---|---|---|---|---|
| | Diff. | p adj | Est. | p adj | Diff. | p adj | Est. | p adj | Est. | p adj | Est. | p adj |
| DMPP- Control | 39.87 | 0.73 | **1.67** | **<0.01** | -0.21 | 0.91 | 3.17 | 0.18 | 0.22 | 0.94 | 2.43 | 0.94 |
| Limonene- Control | 16.84 | 0.99 | -0.50 | 0.68 | -0.06 | 0.99 | 0.84 | 0.98 | 0.13 | 0.99 | 6.45 | 0.90 |
| MBOA- Control | 68.22 | 0.18 | **-1.56** | **0.08** | 0.48 | 0.20 | 1.79 | 0.75 | 0.62 | 0.17 | 6.71 | 0.09 |
| MHPA- Control | **103.85** | **0.00** | **-1.37** | **0.03** | 0.47 | 0.23 | 3.92 | 0.05 | **0.78** | **0.04** | 7.04 | 0.07 |
| Mix- Control | 79.80 | 0.07 | **-3.56** | **<0.01** | -0.41 | 0.37 | **4.05** | **0.04** | 0.67 | 0.11 | 4.58 | 0.05 |
| Limonene - DMPP | -23.02 | 0.96 | **-2.18** | **<0.01** | 0.14 | 0.97 | -2.32 | 0.50 | -0.09 | 0.99 | 8.59 | 0.41 |
| MBOA- DMPP | 28.34 | 0.91 | **-3.23** | **<0.01** | **0.69** | **0.02** | -1.38 | 0.90 | 0.39 | 0.64 | **8.86** | **0.01** |
| MHPA- DMPP | 63.98 | 0.24 | **-3.04** | **<0.01** | **0.68** | **0.02** | 0.74 | 0.99 | 0.55 | 0.28 | **9.19** | **<0.01** |
| Mix- DMPP | 39.93 | 0.73 | **-5.23** | **<0.01** | -0.20 | 0.92 | 0.87 | 0.98 | 0.45 | 0.51 | **4.01** | **<0.01** |
| MBOA- Limonene | 51.37 | 0.48 | -1.05 | 0.43 | 0.54 | 0.11 | 0.94 | 0.97 | 0.49 | 0.41 | 4.27 | 0.55 |
| MHPA- Limonene | **87.01** | **0.04** | -0.86 | 0.43 | 0.53 | 0.12 | 3.07 | 0.20 | 0.65 | 0.14 | 4.60 | 0.48 |
| Mix- Limonene | 62.96 | 0.26 | **-3.06** | **<0.01** | -0.34 | 0.55 | 3.20 | 0.17 | 0.54 | 0.30 | 0.26 | 0.40 |
| MHPA- MBOA | 35.63 | 0.81 | 0.18 | 0.99 | -0.01 | 0.99 | 2.12 | 0.59 | 0.16 | 0.98 | 0.59 | 0.99 |
| Mix- MBOA | 11.58 | 0.99 | **-2.00** | **0.07** | **-0.89** | **<0.01** | 2.25 | 0.53 | 0.05 | 0.99 | 0.33 | 0.99 |
| Mix- MHPA | -24.04 | 0.95 | **-2.19** | **0.02** | **-0.88** | **<0.01** | 0.13 | 0.99 | -0.10 | 0.99 | 2.43 | 0.99 |

**B.**

| Comparison | Total N | | NO2- | | NO3-- | | NH4+ | | DIN | | DON | |
|---|---|---|---|---|---|---|---|---|---|---|---|---|
| | Diff. | p adj | Diff. | p adj | Diff. | p adj | Est. | p adj | Est. | p adj | Est. | p adj |
| DMPP- Control | 20.01 | 0.97 | **-2.23** | **<0.01** | -8.11 | 0.92 | 2.86 | 0.35 | 0.26 | 0.91 | -5.53 | 0.45 |
| Limonene- Control | 19.89 | 0.97 | 1.27 | 0.22 | -9.41 | 0.86 | 1.90 | 0.76 | 0.13 | 0.83 | 2.43 | 0.96 |
| MBOA- Control | 8.97 | 0.99 | 1.09 | 0.38 | 10.28 | 0.81 | -0.37 | 0.99 | 0.10 | 0.98 | 3.23 | 0.88 |
| MHPA- Control | 5.57 | 0.99 | 0.11 | 0.99 | 1.41 | 0.99 | 0.44 | 0.99 | 0.58 | 0.99 | -1.66 | 0.99 |
| Mix- Control | 81.27 | 0.06 | **3.13** | **<0.01** | -15.93 | 0.40 | 4.06 | 0.06 | 0.04 | 0.11 | 3.66 | 0.82 |
| Limonene - DMPP | -0.11 | 1.00 | **3.51** | **<0.01** | -1.29 | 0.99 | -0.95 | 0.98 | -0.08 | 0.99 | 7.97 | 0.10 |
| MBOA- DMPP | -11.04 | 0.99 | **3.32** | **<0.01** | 18.39 | 0.24 | -3.24 | 0.22 | -0.11 | 0.99 | 8.77 | 0.05 |
| MHPA- DMPP | -14.44 | 0.99 | **2.35** | **<0.01** | 9.53 | 0.85 | -2.41 | 0.54 | 0.35 | 0.99 | 3.87 | 0.78 |
| Mix- DMPP | 61.26 | 0.26 | **5.36** | **<0.01** | -7.81 | 0.93 | 1.19 | 0.95 | -0.12 | 0.59 | **9.20** | **0.04** |
| MBOA- Limonene | -10.92 | 0.99 | -0.18 | 0.99 | 19.69 | 0.18 | -2.28 | 0.60 | -0.15 | 0.99 | 0.80 | 0.99 |
| MHPA- Limonene | -14.32 | 0.99 | -1.16 | 0.31 | 10.83 | 0.77 | -1.45 | 0.90 | 0.31 | 0.98 | -4.09 | 0.74 |
| Mix- Limonene | 61.37 | 0.26 | **1.85** | **0.02** | -6.51 | 0.96 | 2.15 | 0.65 | -0.02 | 0.71 | 1.23 | 0.99 |
| MHPA- MBOA | -3.40 | 0.99 | -0.97 | 0.51 | -8.86 | 0.89 | 0.82 | 0.99 | 0.44 | 0.99 | -4.89 | 0.58 |
| Mix- MBOA | 72.30 | 0.12 | **2.04** | **<0.01** | **-26.21** | **0.03** | **4.44** | **0.03** | 0.47 | 0.36 | 0.42 | 0.99 |
| Mix- MHPA | 75.70 | 0.09 | **3.01** | **<0.01** | -17.35 | 0.30 | 3.61 | 0.13 | 0.26 | 0.29 | 5.32 | 0.49 |

**Table S9:** Post-hoc test results for differences in microbial carbon- related processes (Microbial biomass carbon (MBC), microbial respiration (Mic.Resp), microbial growth (Mic.Growth) and carbon use efficiency (CUE) in the **A.** planted and **B.** bare-soil compartment. In all cases, Tukey HSD comparisons were performed following a one-way ANOVA to test for significant differences within each compartment. Significant differences relative to the control and between treatments are shown in bold.

**A.**

| Comparison | MBC | | Mic.Resp | | Mic.Growth | | CUE | |
|---|---|---|---|---|---|---|---|---|
| | Diff. | p adj | Est. | p adj | Diff. | p adj | Est. | p adj |
| DMPP- Control | 0.24 | 0.87 | **-0.99** | **<0.01** | 0.42 | 0.66 | **1.31** | **<0.01** |
| Limonene- Control | -0.11 | 0.99 | -0.21 | 0.87 | -0.13 | 0.99 | -0.01 | 1.00 |
| MBOA- Control | -0.03 | 0.99 | -0.39 | 0.35 | 0.19 | 0.98 | 0.40 | 0.84 |
| MHPA- Control | 0.35 | 0.62 | -0.47 | 0.17 | 0.42 | 0.65 | 0.67 | 0.39 |
| Mix- Control | -0.25 | 0.87 | -0.23 | 0.84 | -0.09 | 0.99 | <0.01 | 1.00 |
| Limonene - DMPP | -0.36 | 0.58 | **0.77** | **<0.01** | -0.55 | 0.37 | **-1.33** | **<0.01** |
| MBOA- DMPP | -0.28 | 0.81 | **0.59** | **0.04** | -0.22 | 0.96 | -0.90 | 0.11 |
| MHPA- DMPP | 0.10 | 0.99 | 0.52 | 0.10 | <0.01 | 1.00 | -0.64 | 0.45 |
| Mix- DMPP | -0.50 | 0.25 | **0.76** | **<0.01** | -0.51 | 0.45 | **-1.32** | **<0.01** |
| MBOA- Limonene | 0.08 | 0.99 | -0.17 | 0.94 | 0.32 | 0.85 | 0.42 | 0.82 |
| MHPA- Limonene | 0.47 | 0.30 | -0.25 | 0.78 | 0.55 | 0.37 | 0.69 | 0.36 |
| Mix- Limonene | -0.13 | 0.99 | -0.01 | 0.99 | 0.03 | 0.99 | 0.01 | 1.00 |
| MHPA- MBOA | 0.38 | 0.52 | -0.07 | 0.99 | 0.22 | 0.96 | 0.26 | 0.97 |
| Mix- MBOA | -0.21 | 0.92 | 0.16 | 0.96 | -0.28 | 0.90 | -0.41 | 0.84 |
| Mix- MHPA | -0.60 | 0.09 | 0.23 | 0.82 | -0.51 | 0.44 | -0.67 | 0.38 |

**B.**

| Comparison | MBC | | Mic.Resp | | Mic.Growth | | CUE | |
|---|---|---|---|---|---|---|---|---|
| | Diff. | p adj | Est. | p adj | Diff. | p adj | Est. | p adj |
| DMPP- Control | 50.87 | 0.96 | **-0.81** | **<0.01** | 0.12 | 0.99 | 0.77 | 0.32 |
| Limonene- Control | -8.99 | 0.99 | 0.43 | 0.33 | -0.14 | 0.99 | -0.52 | 0.72 |
| MBOA- Control | -87.02 | 0.74 | -0.31 | 0.67 | -0.57 | 0.50 | -0.25 | 0.98 |
| MHPA- Control | 34.57 | 0.99 | -0.01 | 0.99 | 0.07 | 0.99 | 0.07 | 0.99 |
| Mix- Control | -7.47 | 0.99 | 0.25 | 0.82 | 0.07 | 0.99 | -0.15 | 0.99 |
| Limonene - DMPP | -59.86 | 0.93 | **1.24** | **<0.01** | -0.27 | 0.95 | **-1.30** | **0.01** |
| MBOA- DMPP | -137.89 | 0.27 | 0.50 | 0.19 | -0.70 | 0.28 | -1.02 | 0.08 |
| MHPA- DMPP | -16.30 | 0.99 | **0.80** | **<0.01** | -0.05 | 0.99 | -0.69 | 0.44 |
| Mix- DMPP | -58.35 | 0.93 | **1.07** | **<0.01** | -0.05 | 0.99 | -0.93 | 0.14 |
| MBOA- Limonene | -78.03 | 0.82 | **-0.74** | **0.01** | -0.42 | 0.77 | 0.27 | 0.97 |

| | | | | | | | | |
|---|---|---|---|---|---|---|---|---|
| MHPA- Limonene | 43.56 | 0.98 | -0.44 | 0.30 | 0.22 | 0.98 | 0.60 | 0.58 |
| Mix- Limonene | 1.51 | 1.00 | -0.17 | 0.96 | 0.21 | 0.98 | 0.36 | 0.92 |
| MHPA- MBOA | 121.59 | 0.40 | 0.30 | 0.70 | 0.64 | 0.36 | 0.33 | 0.94 |
| Mix- MBOA | 79.54 | 0.80 | 0.57 | 0.09 | 0.64 | 0.36 | 0.09 | 0.99 |
| Mix- MHPA | -42.05 | 0.98 | 0.27 | 0.79 | <0.01 | 1.00 | -0.23 | 0.98 |

**Table S10:** Post-hoc test results for differences in $NH_4^+$ concentration in the leachates among treatments within the **A.** planted and **B.** bare-soil compartments. Comparisons between compartments within each treatment are shown in **C**. Pairwise comparisons of estimated marginal means (EMMs) from a gamma–log generalized linear model (GLM) were performed. Significant differences relative to the control and between treatments are shown in bold.

**A**

| **Comparison-** | **Day 0** | | **Day 5** | | **Day 9** | | **Day 14** | |
|---|---|---|---|---|---|---|---|---|
| | **ratio** | **p adj** | **ratio** | **p adj** | **ratio** | **p adj** | **ratio** | **p adj** |
| DMPP-Control | 0.956 | 1.000 | 0.429 | 0.192 | 0.769 | 0.985 | 0.588 | 0.690 |
| Limonene-Control | 1.811 | 0.579 | 1.706 | 0.686 | 1.012 | 1.000 | **0.216** | **0.001** |
| MBOA-Control | 1.057 | 1.000 | **0.224** | **0.001** | 0.525 | 0.487 | **0.285** | **0.010** |
| MHPA-Control | 0.984 | 1.000 | 0.416 | 0.160 | 0.935 | 1.000 | 0.961 | 1.000 |
| Mix-Control | 1.089 | 1.000 | **0.224** | **0.001** | **0.243** | **0.002** | **0.036** | **<0.001** |
| Limonene-DMPP | 1.895 | 0.498 | **3.974** | **0.003** | 1.315 | 0.982 | 0.367 | 0.071 |
| MBOA-DMPP | 1.106 | 1.000 | 0.521 | 0.475 | 0.682 | 0.926 | 0.485 | 0.353 |
| MHPA-DMPP | 1.030 | 1.000 | 0.968 | 1.000 | 1.215 | 0.996 | 1.635 | 0.756 |
| Mix-DMPP | 1.140 | 0.999 | 0.523 | 0.480 | **0.316** | **0.045** | **0.062** | **<0.001** |
| MBOA-Limonene | 0.584 | 0.678 | **0.131** | **<0.001** | 0.519 | 0.467 | 1.321 | 0.973 |
| MHPA-Limonene | 0.544 | 0.551 | **0.244** | **0.002** | 0.924 | 1.000 | **4.458** | **0.001** |
| Mix-Limonene | 0.602 | 0.729 | **0.132** | **<0.001** | **0.240** | **0.002** | **0.168** | **<0.001** |
| MHPA-MBOA | 0.931 | 1.000 | 1.859 | 0.532 | 1.781 | 0.609 | **3.375** | **0.014** |
| Mix-MBOA | 1.031 | 1.000 | 1.003 | 1.000 | 0.463 | 0.286 | **0.127** | **<0.001** |
| Mix-MHPA | 1.107 | 1.000 | 0.540 | 0.538 | 0.260 | 0.004 | **0.038** | **<0.001** |

**B.**

| **Comparison** | **Day 0** | | **Day 5** | | **Day 9** | | **Day 14** | |
|---|---|---|---|---|---|---|---|---|
| | **ratio** | **p adj** | **ratio** | **p adj** | **ratio** | **p adj** | **ratio** | **p adj** |
| DMPP-Control | 0.898 | 1.000 | 0.668 | 0.945 | 0.721 | 0.961 | 1.668 | 0.723 |
| Limonene-Control | 3.213 | 0.598 | 2.874 | 0.175 | 0.974 | 1.000 | 1.946 | 0.451 |
| MBOA-Control | 1.020 | 1.000 | 0.345 | 0.169 | 0.530 | 0.507 | 0.859 | 0.998 |
| MHPA-Control | 0.893 | 1.000 | 0.693 | 0.963 | 0.394 | 0.116 | 2.004 | 0.400 |
| Mix-Control | 1.017 | 1.000 | 0.320 | 0.116 | 0.193 | 0.000 | 0.224 | 0.001 |
| Limonene-DMPP | 3.578 | 0.501 | 4.299 | 0.001 | 1.351 | 0.973 | 1.166 | 0.998 |
| MBOA-DMPP | 1.136 | 1.000 | 0.516 | 0.459 | 0.736 | 0.970 | 0.515 | 0.454 |
| MHPA-DMPP | 0.995 | 1.000 | 1.037 | 1.000 | 0.547 | 0.643 | 1.202 | 0.996 |
| Mix-DMPP | 1.132 | 1.000 | 0.479 | 0.336 | 0.268 | 0.013 | 0.134 | 0.000 |
| MBOA-Limonene | 0.318 | 0.457 | 0.120 | 0.000 | 0.545 | 0.554 | 0.441 | 0.224 |

| | | | | | | | | |
|---|---|---|---|---|---|---|---|---|
| MHPA-Limonene | 0.278 | 0.234 | 0.241 | 0.002 | 0.405 | 0.137 | 1.030 | 1.000 |
| Mix-Limonene | 0.317 | 0.349 | 0.111 | 0.000 | 0.198 | 0.000 | 0.115 | 0.000 |
| MHPA-MBOA | 0.876 | 1.000 | 2.009 | 0.397 | 0.744 | 0.964 | 2.334 | 0.190 |
| Mix-MBOA | 0.997 | 1.000 | 0.928 | 1.000 | 0.364 | 0.068 | 0.260 | 0.004 |
| Mix-MHPA | 1.138 | 0.999 | 0.462 | 0.282 | 0.490 | 0.371 | 0.112 | 0.000 |

**C.**

| **Comparison** | **Day 0** | | **Day 5** | | **Day 9** | | **Day 14** | |
|---|---|---|---|---|---|---|---|---|
| | **ratio** | **p adj** | **ratio** | **p adj** | **ratio** | **p adj** | **ratio** | **p adj** |
| Control Bare- Planted | 0.582 | 0.348 | 0.961 | 0.930 | 1.025 | 0.946 | **2.264** | **0.026** |
| DMPP Bare- Planted | 0.619 | 0.406 | 0.617 | 0.188 | 1.094 | 0.832 | 0.798 | 0.536 |
| Limonene Bare- Planted | 0.328 | 0.055 | 0.571 | 0.126 | 1.065 | 0.864 | **0.251** | **0.000** |
| MBOA Bare- Planted | 0.603 | 0.258 | 0.623 | 0.196 | 1.014 | 0.970 | 0.751 | 0.432 |
| MHPA Bare- Planted | 0.641 | 0.224 | 0.576 | 0.133 | **2.429** | **0.016** | 1.086 | 0.822 |
| Mix Bare- Planted | 0.623 | 0.196 | 0.674 | 0.280 | 1.289 | 0.487 | **0.367** | **0.007** |

**Table S11:** Post-hoc test results for differences in $NO_2^-$ concentration in the leachates among treatments within the **A.** planted and **B.** bare-soil compartments. Comparisons between compartments within each treatment are shown in **C**. Pairwise comparisons of estimated marginal means (EMMs) from a gamma–log generalized linear model (GLM) were performed. Significant differences relative to the control and between treatments are shown in bold**.**

**A.**

| **Comparison** | **Day 0** | | **Day 5** | | **Day 9** | | **Day 14** | |
|---|---|---|---|---|---|---|---|---|
| | **ratio** | **p adj** | **ratio** | **p adj** | **ratio** | **p adj** | **ratio** | **p adj** |
| DMPP-Control | 1.288 | 0.987 | **11.540** | **<0.001** | 1.924 | 0.638 | **4.849** | **0.001** |
| Limonene-Control | **3.432** | **0.025** | 2.354 | 0.254 | **0.227** | **0.003** | **0.248** | **0.007** |
| MBOA-Control | **14.821** | **<0.001** | 0.727 | 0.965 | **0.066** | **<0.001** | **0.203** | **0.001** |
| MHPA-Control | **15.031** | **<0.001** | 1.449 | 0.934 | 0.370 | 0.123 | **4.526** | **0.003** |
| Mix-Control | **17.532** | **<0.001** | **3.867** | **0.010** | **0.080** | **<0.001** | **0.095** | **<0.001** |
| Limonene-DMPP | 2.665 | 0.133 | **0.204** | **0.001** | **0.118** | **<0.001** | **0.051** | **<0.001** |
| MBOA-DMPP | **11.508** | **<0.001** | **0.063** | **<0.001** | **0.034** | **<0.001** | **0.042** | **<0.001** |
| MHPA-DMPP | **11.671** | **<0.001** | **0.126** | **<0.001** | **0.192** | **0.002** | 0.933 | 1.000 |
| Mix-DMPP | **13.614** | **<0.001** | 0.335 | 0.067 | **0.041** | **<0.001** | **0.020** | **<0.001** |
| MBOA-Limonene | **4.318** | **0.004** | **0.309** | **0.038** | **0.289** | **0.024** | 0.822 | 0.996 |
| MHPA-Limonene | **4.379** | **0.003** | 0.616 | 0.819 | 1.634 | 0.811 | **18.277** | **<0.001** |
| Mix-Limonene | **5.108** | **0.001** | 1.643 | 0.804 | 0.352 | 0.090 | 0.385 | 0.153 |
| MHPA-MBOA | 1.014 | 1.000 | 1.994 | 0.497 | **5.645** | **<0.001** | **22.246** | **<0.001** |
| Mix-MBOA | 1.183 | 0.998 | **5.319** | **0.001** | 1.215 | 0.996 | 0.468 | 0.387 |
| Mix-MHPA | 1.166 | 0.999 | 2.668 | 0.132 | **0.215** | **0.002** | **0.021** | **<0.001** |

**B.**

| **Comparison** | **Day 0** | | **Day 5** | | **Day 9** | | **Day 14** | |
|---|---|---|---|---|---|---|---|---|
| | **ratio** | **p adj** | **ratio** | **p adj** | **ratio** | **p adj** | **ratio** | **p adj** |
| DMPP-Control | 0.964 | 1.000 | **5.403** | **0.008** | 2.193 | 0.437 | **12.620** | **<0.001** |
| Limonene-Control | 5.651 | 0.243 | **5.347** | **0.009** | **0.277** | **0.017** | 2.471 | 0.200 |
| MBOA-Control | **8.952** | **0.020** | 0.989 | 1.000 | **0.058** | **<0.001** | 0.735 | 0.970 |
| MHPA-Control | **11.890** | **0.002** | 1.487 | 0.962 | **0.293** | **0.026** | **12.463** | **<0.001** |
| Mix-Control | **12.861** | **0.001** | **4.520** | **0.025** | **0.054** | **<0.001** | **0.196** | **0.001** |
| Limonene-DMPP | 5.859 | 0.222 | 0.990 | 1.000 | **0.126** | **<0.001** | **0.196** | **0.001** |
| MBOA-DMPP | **9.282** | **0.017** | **0.183** | **<0.001** | **0.027** | **<0.001** | **0.058** | **<0.001** |
| MHPA-DMPP | **12.328** | **0.001** | **0.275** | **0.016** | **0.134** | **<0.001** | 0.988 | 1.000 |
| Mix-DMPP | **13.335** | **0.001** | 0.837 | 0.998 | **0.024** | **<0.001** | **0.016** | **<0.001** |
| MBOA-Limonene | 1.584 | 0.984 | **0.185** | **<0.001** | **0.211** | **0.002** | **0.298** | **0.029** |

| | | | | | | | | |
|---|---|---|---|---|---|---|---|---|
| MHPA-Limonene | 2.104 | 0.837 | **0.278** | **0.018** | 1.061 | 1.000 | **5.044** | **0.001** |
| Mix-Limonene | 2.276 | 0.771 | 0.845 | 0.998 | **0.194** | **0.001** | **0.079** | **<0.001** |
| MHPA-MBOA | 1.328 | 0.992 | 1.503 | 0.904 | **5.027** | **0.001** | **16.953** | **<0.001** |
| Mix-MBOA | 1.437 | 0.975 | **4.569** | **0.002** | 0.918 | 1.000 | **0.267** | **0.013** |
| Mix-MHPA | 1.082 | 1.000 | 3.040 | 0.059 | **0.183** | **<0.001** | **0.016** | **<0.001** |

**C.**

| **Comparison** | **Day 0** | | **Day 5** | | **Day 9** | | **Day 14** | |
|---|---|---|---|---|---|---|---|---|
| | **ratio** | **p adj** | **ratio** | **p adj** | **ratio** | **p adj** | **ratio** | **p adj** |
| Control Bare- Planted | 0.734 | 0.619 | 0.773 | 0.593 | 1.140 | 0.739 | 2.603 | 0.016 |
| DMPP Bare- Planted | 0.980 | 0.973 | 1.650 | 0.204 | 1.000 | 1.000 | 1.000 | 1.000 |
| Limonene Bare- Planted | 0.446 | 0.195 | **0.340** | **0.007** | 0.934 | 0.861 | **0.261** | **0.001** |
| MBOA Bare- Planted | 1.214 | 0.687 | 0.568 | 0.152 | 1.280 | 0.530 | 0.720 | 0.405 |
| MHPA Bare- Planted | 0.927 | 0.848 | 0.753 | 0.472 | 1.438 | 0.357 | 0.945 | 0.886 |
| Mix Bare- Planted | 1.000 | 1.000 | 0.661 | 0.294 | 1.694 | 0.182 | 1.262 | 0.554 |

**Table S12:** Post-hoc test results for differences in $NO_3^-$ concentration in the leachates among treatments within the **A.** planted and **B.** bare-soil compartments. Comparisons between compartments within each treatment are shown in **C**. Pairwise comparisons of estimated marginal means (EMMs) from a gamma–log generalized linear model (GLM) were performed. Significant differences relative to the control and between treatments are shown in bold.

**A.**

| Comparison | Day 0 | | Day 5 | | Day 9 | | Day 14 | |
|---|---|---|---|---|---|---|---|---|
| | **ratio** | **p adj** | **ratio** | **p adj** | **ratio** | **p adj** | **ratio** | **p adj** |
| DMPP-Control | 0.831 | 0.997 | 1.052 | 1.000 | 1.717 | 0.776 | 2.867 | 0.069 |
| Limonene-Control | 1.421 | 0.940 | 1.033 | 1.000 | 1.487 | 0.903 | 0.585 | 0.721 |
| MBOA-Control | 1.425 | 0.938 | 1.967 | 0.484 | **3.437** | **0.018** | 1.760 | 0.675 |
| MHPA-Control | 1.270 | 0.989 | 1.379 | 0.958 | 1.938 | 0.509 | 2.174 | 0.325 |
| Mix-Control | 1.107 | 1.000 | 2.544 | 0.146 | **4.576** | **0.001** | 1.546 | 0.862 |
| Limonene-DMPP | 1.711 | 0.721 | 0.982 | 1.000 | 0.866 | 0.999 | **0.204** | **0.001** |
| MBOA-DMPP | 1.716 | 0.716 | 1.869 | 0.572 | 2.002 | 0.542 | 0.614 | 0.794 |
| MHPA-DMPP | 1.528 | 0.875 | 1.311 | 0.980 | 1.129 | 1.000 | 0.758 | 0.978 |
| Mix-DMPP | 1.332 | 0.975 | 2.417 | 0.194 | 2.665 | 0.170 | 0.539 | 0.585 |
| MBOA-Limonene | 1.003 | 1.000 | 1.903 | 0.540 | 2.311 | 0.244 | 3.009 | 0.050 |
| MHPA-Limonene | 0.894 | 1.000 | 1.335 | 0.974 | 1.304 | 0.982 | **3.719** | **0.010** |
| Mix-Limonene | 0.779 | 0.986 | 2.462 | 0.176 | **3.077** | **0.042** | 2.644 | 0.116 |
| MHPA-MBOA | 0.891 | 1.000 | 0.701 | 0.937 | 0.564 | 0.662 | 1.236 | 0.994 |
| Mix-MBOA | 0.776 | 0.985 | 1.293 | 0.984 | 1.331 | 0.975 | 0.879 | 0.999 |
| Mix-MHPA | 0.871 | 0.999 | 1.844 | 0.595 | 2.361 | 0.220 | 0.711 | 0.947 |

**B.**

| Comparison | Day 0 | | Day 5 | | Day 9 | | Day 14 | |
|---|---|---|---|---|---|---|---|---|
| | **ratio** | **p adj** | **ratio** | **p adj** | **ratio** | **p adj** | **ratio** | **p adj** |
| DMPP-Control | 0.678 | 0.996 | 1.206 | 0.999 | 2.760 | 0.141 | **4.848** | **0.001** |
| Limonene-Control | 2.033 | 0.938 | 2.194 | 0.544 | 1.258 | 0.991 | 1.489 | 0.901 |
| MBOA-Control | 2.499 | 0.733 | 3.210 | 0.131 | 2.795 | 0.082 | 1.799 | 0.637 |
| MHPA-Control | 2.063 | 0.835 | 2.942 | 0.196 | 1.331 | 0.975 | 2.500 | 0.161 |
| Mix-Control | 1.295 | 0.998 | 3.681 | 0.065 | 3.149 | 0.036 | 1.625 | 0.798 |
| Limonene-DMPP | 3.000 | 0.701 | 1.819 | 0.618 | 0.456 | 0.399 | **0.307** | **0.028** |
| MBOA-DMPP | 3.687 | 0.360 | 2.662 | 0.112 | 1.013 | 1.000 | 0.371 | 0.103 |
| MHPA-DMPP | 3.044 | 0.438 | 2.440 | 0.185 | 0.482 | 0.486 | 0.516 | 0.508 |
| Mix-DMPP | 1.911 | 0.890 | **3.052** | **0.045** | 1.141 | 1.000 | 0.335 | 0.053 |
| MBOA-Limonene | 1.229 | 1.000 | 1.464 | 0.917 | 2.223 | 0.294 | 1.208 | 0.996 |

| | | | | | | | | |
|---|---|---|---|---|---|---|---|---|
| MHPA-Limonene | 1.015 | 1.000 | 1.341 | 0.972 | 1.058 | 1.000 | 1.679 | 0.750 |
| Mix-Limonene | 0.637 | 0.975 | 1.678 | 0.751 | 2.504 | 0.160 | 1.091 | 1.000 |
| MHPA-MBOA | 0.826 | 0.998 | 0.916 | 1.000 | 0.476 | 0.377 | 1.390 | 0.954 |
| Mix-MBOA | 0.518 | 0.721 | 1.146 | 0.999 | 1.127 | 1.000 | 0.903 | 1.000 |
| Mix-MHPA | 0.628 | 0.825 | 1.251 | 0.992 | 2.366 | 0.217 | 0.650 | 0.867 |

**C.**

| Comparison | Day 0 | | Day 5 | | Day 9 | | Day 14 | |
|---|---|---|---|---|---|---|---|---|
| | **ratio** | **p adj** | **ratio** | **p adj** | **ratio** | **p adj** | **ratio** | **p adj** |
| Control Bare- Planted | 0.660 | 0.492 | 1.167 | 0.741 | 1.273 | 0.528 | 1.494 | 0.294 |
| DMPP Bare- Planted | 0.810 | 0.726 | 1.019 | 0.962 | 0.792 | 0.596 | 0.883 | 0.745 |
| Limonene Bare- Planted | 0.462 | 0.201 | 0.550 | 0.118 | 1.505 | 0.285 | 0.587 | 0.164 |
| MBOA Bare- Planted | **0.377** | **0.038** | 0.715 | 0.380 | 1.565 | 0.242 | 1.461 | 0.321 |
| MHPA Bare- Planted | **0.407** | **0.020** | 0.547 | 0.115 | 1.853 | 0.107 | 1.299 | 0.493 |
| Mix Bare- Planted | 0.564 | 0.135 | 0.807 | 0.573 | 1.849 | 0.109 | 1.421 | 0.357 |

**Table S13:** Post-hoc test results for differences in $N_2O$ concentration in the planted compartment during the **A.** first (day 0) **B.** second (day 5), **C.** third (day 9), and **B.** fourth (day 14) watering event. Pairwise comparisons of estimated marginal means (EMMs) from a gamma–log generalized linear model (GLM) were performed. Significant differences relative to the control and between treatments are shown in bold.

**A.**

| Comparison | 0h | | 7h | | 24h | |
|---|---|---|---|---|---|---|
| | ratio | p adj | ratio | p adj | ratio | p adj |
| DMPP-Control | 1.671 | 0.908 | 3.120 | 0.208 | 1.668 | 0.909 |
| Limonene-Control | 1.925 | 0.779 | 1.933 | 0.775 | 1.216 | 0.999 |
| MBOA-Control | 1.500 | 0.965 | 1.881 | 0.804 | 1.562 | 0.948 |
| MHPA-Control | 1.548 | 0.952 | 1.677 | 0.906 | 0.621 | 0.932 |
| Mix-Control | 2.057 | 0.701 | **4.618** | **0.030** | 3.921 | 0.073 |
| Limonene-DMPP | 1.151 | 1.000 | 0.620 | 0.931 | 0.729 | 0.988 |
| MBOA-DMPP | 0.898 | 1.000 | 0.603 | 0.914 | 0.936 | 1.000 |
| MHPA-DMPP | 0.926 | 1.000 | 0.538 | 0.816 | 0.372 | 0.358 |
| Mix-DMPP | 1.230 | 0.998 | 1.480 | 0.970 | 2.350 | 0.527 |
| MBOA-Limonene | 0.779 | 0.996 | 0.973 | 1.000 | 1.284 | 0.996 |
| MHPA-Limonene | 0.804 | 0.998 | 0.868 | 1.000 | 0.510 | 0.759 |
| Mix-Limonene | 1.069 | 1.000 | 2.390 | 0.505 | 3.224 | 0.182 |
| MHPA-MBOA | 1.032 | 1.000 | 0.892 | 1.000 | 0.398 | 0.439 |
| Mix-MBOA | 1.371 | 0.989 | 2.455 | 0.470 | 2.511 | 0.441 |
| Mix-MHPA | 1.329 | 0.993 | 2.753 | 0.331 | **6.317** | **0.004** |

**B.**

| Comparison | 0h | | 7h | | 24h | |
|---|---|---|---|---|---|---|
| | ratio | p adj | ratio | p adj | ratio | p adj |
| DMPP-Control | 1.739 | 0.878 | 2.848 | 0.294 | **6.343** | **0.004** |
| Limonene-Control | 1.607 | 0.933 | **0.090** | **<0.001** | 0.432 | 0.548 |
| MBOA-Control | 1.146 | 1.000 | **0.121** | **0.001** | 0.343 | 0.271 |
| MHPA-Control | 1.162 | 1.000 | **0.097** | **<0.001** | **0.216** | **0.029** |
| Mix-Control | 1.823 | 0.836 | **0.156** | **0.004** | 0.317 | 0.199 |
| Limonene-DMPP | 0.924 | 1.000 | **0.032** | **<0.001** | **0.068** | **<0.001** |
| MBOA-DMPP | 0.659 | 0.961 | **0.043** | **<0.001** | **0.054** | **<0.001** |
| MHPA-DMPP | 0.668 | 0.966 | **0.034** | **<0.001** | **0.034** | **<0.001** |
| Mix-DMPP | 1.048 | 1.000 | **0.055** | **<0.001** | **0.050** | **<0.001** |
| MBOA-Limonene | 0.713 | 0.984 | 1.346 | 0.991 | 0.794 | 0.997 |
| MHPA-Limonene | 0.723 | 0.987 | 1.080 | 1.000 | 0.499 | 0.734 |

| | | | | | | |
|---|---|---|---|---|---|---|
| Mix-Limonene | 1.135 | 1.000 | 1.738 | 0.879 | 0.734 | 0.989 |
| MHPA-MBOA | 1.013 | 1.000 | 0.803 | 0.998 | 0.629 | 0.939 |
| Mix-MBOA | 1.591 | 0.939 | 1.291 | 0.996 | 0.924 | 1.000 |
| Mix-MHPA | 1.570 | 0.946 | 1.609 | 0.933 | 1.468 | 0.972 |

**C.**

| Comparison | 0h | | 7h | | 24h | |
|---|---|---|---|---|---|---|
| | ratio | p adj | ratio | p adj | ratio | p adj |
| DMPP-Control | 1.797 | 0.850 | 1.637 | 0.922 | 1.741 | 0.877 |
| Limonene-Control | 1.955 | 0.762 | 0.371 | 0.354 | 0.641 | 0.949 |
| MBOA-Control | 1.376 | 0.988 | **0.170** | **0.006** | **0.051** | **<0.001** |
| MHPA-Control | 1.467 | 0.973 | **0.121** | **0.001** | 0.353 | 0.301 |
| Mix-Control | 1.944 | 0.768 | **0.035** | **<0.001** | **0.117** | **<0.001** |
| Limonene-DMPP | 1.088 | 1.000 | **0.226** | **0.038** | 0.368 | 0.347 |
| MBOA-DMPP | 0.766 | 0.995 | **0.104** | **<0.001** | **0.029** | **<0.001** |
| MHPA-DMPP | 0.817 | 0.999 | **0.074** | **<0.001** | **0.203** | **0.020** |
| Mix-DMPP | 1.082 | 1.000 | **0.022** | **<0.001** | **0.067** | **<0.001** |
| MBOA-Limonene | 0.704 | 0.982 | 0.459 | 0.626 | **0.080** | **<0.001** |
| MHPA-Limonene | 0.751 | 0.993 | 0.327 | 0.226 | 0.551 | 0.840 |
| Mix-Limonene | 0.995 | 1.000 | **0.096** | **<0.001** | **0.183** | **0.010** |
| MHPA-MBOA | 1.066 | 1.000 | 0.713 | 0.984 | **6.902** | **0.002** |
| Mix-MBOA | 1.413 | 0.983 | **0.208** | **0.024** | 2.290 | 0.561 |
| Mix-MHPA | 1.325 | 0.993 | 0.292 | 0.141 | 0.332 | 0.239 |

**D.**

| Comparison | 0h | | 7h | | 24h | |
|---|---|---|---|---|---|---|
| | ratio | p adj | ratio | p adj | ratio | p adj |
| DMPP-Control | 1.460 | 0.974 | **4.689** | **0.027** | **5.831** | **0.007** |
| Limonene-Control | 1.516 | 0.961 | 0.635 | 0.944 | 0.983 | 1.000 |
| MBOA-Control | 1.300 | 0.995 | 2.081 | 0.686 | 0.336 | 0.251 |
| MHPA-Control | 1.350 | 0.991 | 2.168 | 0.633 | 0.764 | 0.994 |
| Mix-Control | 1.728 | 0.883 | **0.124** | **0.001** | 0.462 | 0.635 |
| Limonene-DMPP | 1.038 | 1.000 | **0.135** | **0.001** | **0.169** | **0.006** |
| MBOA-DMPP | 0.890 | 1.000 | 0.444 | 0.583 | **0.058** | **<0.001** |
| MHPA-DMPP | 0.924 | 1.000 | 0.462 | 0.637 | **0.131** | **0.001** |
| Mix-DMPP | 1.183 | 0.999 | **0.027** | **<0.001** | **0.079** | **<0.001** |
| MBOA-Limonene | 0.858 | 1.000 | 3.276 | 0.170 | 0.342 | 0.267 |
| MHPA-Limonene | 0.891 | 1.000 | 3.413 | 0.142 | 0.777 | 0.996 |
| Mix-Limonene | 1.140 | 1.000 | **0.196** | **0.016** | 0.469 | 0.656 |
| MHPA-MBOA | 1.038 | 1.000 | 1.042 | 1.000 | 2.271 | 0.572 |
| Mix-MBOA | 1.329 | 0.993 | **0.060** | **<0.001** | 1.373 | 0.988 |
| Mix-MHPA | 1.280 | 0.996 | **0.057** | **<0.001** | 0.604 | 0.915 |

**Table S14:** Post-hoc test results for differences in $N_2O$ concentration in the bare-soil compartment during the **A.** first (day 0) **B.** second (day 5), **C.** third (day 9), and **B.** fourth (day 14) watering event. Pairwise comparisons of estimated marginal means (EMMs) from a generalized least square model (GLS) were performed. Significant differences relative to the control and between treatments are shown in bold.

**A.**

| Comparison | 0h | | 7h | | 24h | |
|---|---|---|---|---|---|---|
| | **Estimate** | **p adj** | **Estimate** | **p adj** | **Estimate** | **p adj** |
| DMPP-Control | <0.001 | 1.000 | 0.002 | 1.000 | -0.002 | 1.000 |
| Limonene-Control | <0.001 | 1.000 | 0.002 | 1.000 | 0.003 | 1.000 |
| MBOA-Control | <0.001 | 1.000 | 0.002 | 1.000 | 0.002 | 1.000 |
| MHPA-Control | <0.001 | 1.000 | 0.001 | 1.000 | 0.001 | 1.000 |
| Mix-Control | <0.001 | 1.000 | 0.003 | 1.000 | <0.001 | 1.000 |
| Limonene-DMPP | <0.001 | 1.000 | <0.001 | 1.000 | 0.005 | 1.000 |
| MBOA-DMPP | <0.001 | 1.000 | <0.001 | 1.000 | 0.004 | 1.000 |
| MHPA-DMPP | <0.001 | 1.000 | <0.001 | 1.000 | 0.003 | 1.000 |
| Mix-DMPP | <0.001 | 1.000 | 0.001 | 1.000 | 0.001 | 1.000 |
| MBOA-Limonene | <0.001 | 1.000 | <0.001 | 1.000 | -0.001 | 1.000 |
| MHPA-Limonene | <0.001 | 1.000 | -0.001 | 1.000 | -0.002 | 1.000 |
| Mix-Limonene | <0.001 | 1.000 | 0.001 | 1.000 | -0.004 | 1.000 |
| MHPA-MBOA | <0.001 | 1.000 | <0.001 | 1.000 | -0.001 | 1.000 |
| Mix-MBOA | <0.001 | 1.000 | 0.002 | 1.000 | -0.002 | 1.000 |
| Mix-MHPA | <0.001 | 1.000 | 0.002 | 1.000 | -0.001 | 1.000 |

**B.**

| Comparison | 0h | | 7h | | 24h | |
|---|---|---|---|---|---|---|
| | **Estimate** | **p adj** | **Estimate** | **p adj** | **Estimate** | **p adj** |
| DMPP-Control | <0.001 | 1.000 | <0.001 | 1.000 | <0.001 | 1.000 |
| Limonene-Control | <0.001 | 1.000 | -0.011 | 0.984 | -0.015 | 0.928 |
| MBOA-Control | <0.001 | 1.000 | -0.005 | 1.000 | -0.004 | 1.000 |
| MHPA-Control | <0.001 | 1.000 | -0.006 | 0.999 | -0.009 | 0.991 |
| Mix-Control | <0.001 | 1.000 | -0.007 | 0.998 | -0.008 | 0.995 |
| Limonene-DMPP | <0.001 | 1.000 | -0.011 | 0.982 | -0.015 | 0.929 |
| MBOA-DMPP | <0.001 | 1.000 | -0.005 | 1.000 | -0.004 | 1.000 |
| MHPA-DMPP | <0.001 | 1.000 | -0.006 | 0.999 | -0.009 | 0.992 |
| Mix-DMPP | <0.001 | 1.000 | -0.007 | 0.998 | -0.008 | 0.996 |
| MBOA-Limonene | <0.001 | 1.000 | 0.006 | 0.999 | 0.011 | 0.981 |
| MHPA-Limonene | <0.001 | 1.000 | 0.005 | 1.000 | 0.006 | 0.999 |

| | | | | | | |
|---|---|---|---|---|---|---|
| Mix-Limonene | <0.001 | 1.000 | 0.004 | 1.000 | 0.007 | 0.998 |
| MHPA-MBOA | <0.001 | 1.000 | -0.002 | 1.000 | -0.005 | 0.999 |
| Mix-MBOA | <0.001 | 1.000 | -0.002 | 1.000 | -0.004 | 1.000 |
| Mix-MHPA | <0.001 | 1.000 | -0.001 | 1.000 | 0.001 | 1.000 |

**C.**

| Comparison | 0h | | 7h | | 24h | |
|---|---|---|---|---|---|---|
| | **Estimate** | **p adj** | **Estimate** | **p adj** | **Estimate** | **p adj** |
| DMPP-Control | <0.001 | 1.000 | <0.001 | 1.000 | <0.001 | 1.000 |
| Limonene-Control | <0.001 | 1.000 | -0.012 | 0.974 | -0.005 | 1.000 |
| MBOA-Control | <0.001 | 1.000 | -0.003 | 1.000 | -0.001 | 1.000 |
| MHPA-Control | <0.001 | 1.000 | -0.001 | 1.000 | -0.005 | 1.000 |
| Mix-Control | <0.001 | 1.000 | -0.023 | 0.714 | -0.004 | 1.000 |
| Limonene-DMPP | <0.001 | 1.000 | -0.012 | 0.977 | -0.004 | 1.000 |
| MBOA-DMPP | <0.001 | 1.000 | -0.002 | 1.000 | -0.001 | 1.000 |
| MHPA-DMPP | <0.001 | 1.000 | -0.001 | 1.000 | -0.004 | 1.000 |
| Mix-DMPP | <0.001 | 1.000 | -0.022 | 0.729 | -0.004 | 1.000 |
| MBOA-Limonene | <0.001 | 1.000 | 0.010 | 0.991 | 0.003 | 1.000 |
| MHPA-Limonene | <0.001 | 1.000 | 0.011 | 0.984 | <0.001 | 1.000 |
| Mix-Limonene | <0.001 | 1.000 | -0.010 | 0.986 | 0.001 | 1.000 |
| MHPA-MBOA | <0.001 | 1.000 | 0.001 | 1.000 | -0.004 | 1.000 |
| Mix-MBOA | <0.001 | 1.000 | -0.020 | 0.806 | -0.003 | 1.000 |
| Mix-MHPA | <0.001 | 1.000 | -0.021 | 0.761 | 0.001 | 1.000 |

**D.**

| Comparison | 0h | | 7h | | 24h | |
|---|---|---|---|---|---|---|
| | **Estimate** | **p adj** | **Estimate** | **p adj** | **Estimate** | **p adj** |
| DMPP-Control | <0.001 | 1.000 | 0.005 | 0.999 | 0.007 | 0.997 |
| Limonene-Control | <0.001 | 1.000 | **-0.104** | **<0.001** | **-0.101** | **<0.001** |
| MBOA-Control | <0.001 | 1.000 | -0.002 | 1.000 | -0.001 | 1.000 |
| MHPA-Control | <0.001 | 1.000 | -0.005 | 0.999 | -0.020 | 0.808 |
| Mix-Control | <0.001 | 1.000 | -0.014 | 0.955 | 0.002 | 1.000 |
| Limonene-DMPP | <0.001 | 1.000 | **-0.109** | **<0.001** | **-0.108** | **<0.001** |
| MBOA-DMPP | <0.001 | 1.000 | -0.007 | 0.998 | -0.008 | 0.995 |
| MHPA-DMPP | <0.001 | 1.000 | -0.011 | 0.985 | -0.027 | 0.522 |
| Mix-DMPP | <0.001 | 1.000 | -0.019 | 0.842 | -0.005 | 1.000 |
| MBOA-Limonene | <0.001 | 1.000 | **0.102** | **<0.001** | **0.100** | **<0.001** |
| MHPA-Limonene | <0.001 | 1.000 | **0.099** | **<0.001** | **0.081** | **<0.001** |
| Mix-Limonene | <0.001 | 1.000 | **0.091** | **<0.001** | **0.103** | **<0.001** |
| MHPA-MBOA | <0.001 | 1.000 | -0.004 | 1.000 | -0.019 | 0.841 |
| Mix-MBOA | <0.001 | 1.000 | -0.012 | 0.975 | 0.003 | 1.000 |
| Mix-MHPA | <0.001 | 1.000 | -0.008 | 0.995 | 0.022 | 0.720 |

**Table S15:** Post-hoc test results for differences in $NO_2^-$ concentration in the pore water among treatments within the **A.** planted and **B.** bare-soil compartments. Pairwise comparisons of estimated marginal means (EMMs) from a gamma–log generalized linear model (GLM) were performed. Significant differences relative to the control and between treatments are shown in bold.

**A.**

| Comparison | Day 0 | | Day 5 | | Day 9 | | Day 14 | |
|---|---|---|---|---|---|---|---|---|
| | ratio | p adj | ratio | p adj | ratio | p adj | ratio | p adj |
| DMPP-Control | **2.226** | **0.014** | **11.858** | **<0.001** | 1.215 | 0.965 | **21.542** | **<0.001** |
| Limonene-Control | **5.792** | **<0.001** | **4.545** | **<0.001** | **0.200** | **<0.001** | 0.754 | 0.930 |
| MBOA-Control | **21.908** | **<0.001** | 1.180 | 0.983 | **0.039** | **<0.001** | 0.859 | 0.988 |
| MHPA-Control | **17.135** | **<0.001** | 1.369 | 0.781 | 0.547 | 0.130 | 0.829 | 0.988 |
| Mix-Control | **25.767** | **<0.001** | **2.701** | **0.001** | **0.054** | **<0.001** | 1.049 | 1.000 |
| Limonene-DMPP | **2.602** | **0.002** | **0.383** | **0.002** | **0.164** | **<0.001** | **0.035** | **<0.001** |
| MBOA-DMPP | **9.843** | **<0.001** | **0.100** | **<0.001** | **0.032** | **<0.001** | **0.040** | **<0.001** |
| MHPA-DMPP | **7.698** | **<0.001** | **0.115** | **<0.001** | **0.450** | **0.015** | **0.039** | **<0.001** |
| Mix-DMPP | **11.577** | **<0.001** | **0.228** | **<0.001** | **0.044** | **<0.001** | **0.049** | **<0.001** |
| MBOA-Limonene | **3.783** | **<0.001** | **0.260** | **<0.001** | **0.195** | **<0.001** | 1.139 | 0.998 |
| MHPA-Limonene | **2.958** | **<0.001** | **0.301** | **<0.001** | **2.740** | **0.001** | 1.100 | 1.000 |
| Mix-Limonene | **4.449** | **<0.001** | 0.594 | 0.262 | **0.270** | **<0.001** | 1.392 | 0.968 |
| MHPA-MBOA | 0.782 | 0.910 | 1.160 | 0.990 | **14.033** | **<0.001** | 0.966 | 1.000 |
| Mix-MBOA | 1.176 | 0.984 | **2.289** | **0.010** | 1.381 | 0.816 | 1.221 | 0.995 |
| Mix-MHPA | 1.504 | 0.537 | 1.973 | 0.060 | **0.098** | **<0.001** | 1.265 | 0.993 |

**B.**

| Comparison | Day 0 | | Day 5 | | Day 9 | | Day 14 | |
|---|---|---|---|---|---|---|---|---|
| | ratio | p adj | ratio | p adj | ratio | p adj | ratio | p adj |
| DMPP-Control | 1.838 | 0.123 | **5.257** | **<0.001** | 1.000 | 1.000 | **16.502** | **<0.001** |
| Limonene-Control | **6.913** | **<0.001** | 1.252 | 0.954 | **0.073** | **<0.001** | 0.883 | 0.995 |
| MBOA-Control | **10.261** | **<0.001** | 1.152 | 0.992 | **0.053** | **<0.001** | 0.960 | 1.000 |
| MHPA-Control | **9.020** | **<0.001** | 1.200 | 0.974 | 0.684 | 0.615 | 1.354 | 0.807 |
| Mix-Control | **17.458** | **<0.001** | 1.283 | 0.904 | **0.064** | **<0.001** | 0.990 | 1.000 |
| Limonene-DMPP | **3.760** | **<0.001** | **0.238** | **<0.001** | **0.073** | **<0.001** | **0.054** | **<0.001** |
| MBOA-DMPP | **5.581** | **<0.001** | **0.219** | **<0.001** | **0.053** | **<0.001** | **0.058** | **<0.001** |
| MHPA-DMPP | **4.906** | **<0.001** | **0.228** | **<0.001** | 0.684 | 0.615 | **0.082** | **<0.001** |
| Mix-DMPP | **9.496** | **<0.001** | **0.244** | **<0.001** | **0.064** | **<0.001** | **0.060** | **<0.001** |
| MBOA-Limonene | 1.484 | 0.572 | 0.920 | 1.000 | 0.715 | 0.730 | 1.087 | 0.999 |
| MHPA-Limonene | 1.305 | 0.909 | 0.959 | 1.000 | **9.310** | **<0.001** | 1.533 | 0.485 |

| | | | | | | | | |
|---|---|---|---|---|---|---|---|---|
| Mix-Limonene | **2.526** | **0.002** | 1.025 | 1.000 | 0.864 | 0.993 | 1.122 | 0.999 |
| MHPA-MBOA | 0.879 | 0.996 | 1.042 | 1.000 | **13.025** | **<0.001** | 1.410 | 0.709 |
| Mix-MBOA | 1.701 | 0.241 | 1.114 | 0.998 | 1.209 | 0.978 | 1.032 | 1.000 |
| Mix-MHPA | 1.936 | 0.120 | 1.069 | 1.000 | **0.093** | **<0.001** | 0.732 | 0.896 |

**Table S16:** Post-hoc test results for differences in $NO_3^-$ concentration in the pore water among treatments within the **A.** planted and **B.** bare-soil compartments. Pairwise comparisons of estimated marginal means (EMMs) from a gamma–log generalized linear model (GLM) were performed. Significant differences relative to the control and between treatments are shown in bold.

**A.**

| Comparison | Day 0 | | Day 5 | | Day 9 | | Day 14 | |
|---|---|---|---|---|---|---|---|---|
| | ratio | p adj | ratio | p adj | ratio | p adj | ratio | p adj |
| DMPP-Control | 0.911 | 0.921 | 0.970 | 0.999 | 0.965 | 0.999 | 1.072 | 0.976 |
| Limonene-Control | 1.043 | 0.998 | 1.265 | 0.159 | **1.428** | **0.003** | 1.101 | 0.933 |
| MBOA-Control | 1.259 | 0.148 | **1.872** | **<0.001** | **1.573** | **<0.001** | 1.185 | 0.467 |
| MHPA-Control | 1.089 | 0.952 | 1.147 | 0.691 | 1.042 | 0.998 | 0.982 | 1.000 |
| Mix-Control | **1.513** | **<0.001** | **2.379** | **<0.001** | **2.573** | **<0.001** | **2.504** | **<0.001** |
| Limonene-DMPP | 1.145 | 0.705 | 1.305 | 0.076 | **1.481** | **0.001** | 1.027 | 1.000 |
| MBOA-DMPP | **1.382** | **0.010** | **1.930** | **<0.001** | **1.631** | **<0.001** | 1.105 | 0.896 |
| MHPA-DMPP | 1.196 | 0.449 | 1.183 | 0.478 | 1.080 | 0.964 | 0.916 | 0.955 |
| Mix-DMPP | **1.661** | **<0.001** | **2.453** | **<0.001** | **2.667** | **<0.001** | **2.336** | **<0.001** |
| MBOA-Limonene | 1.207 | 0.349 | **1.480** | **0.001** | 1.102 | 0.908 | 1.076 | 0.979 |
| MHPA-Limonene | 1.044 | 0.998 | 0.907 | 0.917 | **0.729** | **0.013** | 0.892 | 0.898 |
| Mix-Limonene | **1.451** | **0.002** | **1.880** | **<0.001** | **1.801** | **<0.001** | **2.274** | **<0.001** |
| MHPA-MBOA | 0.865 | 0.675 | **0.613** | **<0.001** | **0.662** | **<0.001** | 0.829 | 0.440 |
| Mix-MBOA | 1.202 | 0.372 | 1.271 | 0.118 | **1.635** | **<0.001** | **2.113** | **<0.001** |
| Mix-MHPA | **1.389** | **0.012** | **2.073** | **<0.001** | **2.470** | **<0.001** | **2.550** | **<0.001** |

**B.**

| Comparison | Day 0 | | Day 5 | | Day 9 | | Day 14 | |
|---|---|---|---|---|---|---|---|---|
| | ratio | p adj | ratio | p adj | ratio | p adj | ratio | p adj |
| DMPP-Control | 0.911 | 0.921 | 0.970 | 0.999 | 0.965 | 0.999 | 1.072 | 0.976 |
| Limonene-Control | 1.043 | 0.998 | 1.265 | 0.159 | **1.428** | **0.003** | 1.101 | 0.933 |
| MBOA-Control | 1.259 | 0.148 | **1.872** | **<0.001** | **1.573** | **<0.001** | 1.185 | 0.467 |
| MHPA-Control | 1.089 | 0.952 | 1.147 | 0.691 | 1.042 | 0.998 | 0.982 | 1.000 |
| Mix-Control | **1.513** | **<0.001** | **2.379** | **<0.001** | **2.573** | **<0.001** | **2.504** | **<0.001** |
| Limonene-DMPP | 1.145 | 0.705 | 1.305 | 0.076 | **1.481** | **0.001** | 1.027 | 1.000 |
| MBOA-DMPP | **1.382** | **0.010** | **1.930** | **<0.001** | **1.631** | **<0.001** | 1.105 | 0.896 |
| MHPA-DMPP | 1.196 | 0.449 | 1.183 | 0.478 | 1.080 | 0.964 | 0.916 | 0.955 |
| Mix-DMPP | **1.661** | **<0.001** | **2.453** | **<0.001** | **2.667** | **<0.001** | **2.336** | **<0.001** |
| MBOA-Limonene | 1.207 | 0.349 | **1.480** | **0.001** | 1.102 | 0.908 | 1.076 | 0.979 |
| MHPA-Limonene | 1.044 | 0.998 | 0.907 | 0.917 | **0.729** | **0.013** | 0.892 | 0.898 |

| | | | | | | | | |
|---|---|---|---|---|---|---|---|---|
| Mix-Limonene | **1.451** | **0.002** | **1.880** | **<0.001** | **1.801** | **<0.001** | **2.274** | **<0.001** |
| MHPA-MBOA | 0.865 | 0.675 | **0.613** | **<0.001** | **0.662** | **<0.001** | 0.829 | 0.440 |
| Mix-MBOA | 1.202 | 0.372 | 1.271 | 0.118 | **1.635** | **<0.001** | **2.113** | **<0.001** |
| Mix-MHPA | **1.389** | **0.012** | **2.073** | **<0.001** | **2.470** | **<0.001** | **2.550** | **<0.001** |

**Table S17:** Post hoc test results for differences in $NO_2^-$ as a fraction of applied N among treatments within the **A.** planted and **B.** bare-soil compartments. Pairwise comparisons of estimated marginal means (EMMs) from a gamma–log generalized linear model (GLM) were performed. Significant differences relative to the control and between treatments are shown in bold.

**A.**

| Comparison | Day 0 | | Day 5 | | Day 9 | | Day 14 | |
|---|---|---|---|---|---|---|---|---|
| | ratio | p adj | ratio | p adj | ratio | p adj | ratio | p adj |
| DMPP-Control | **1.307** | **<0.001** | **1.516** | **<0.001** | 1.006 | 1.000 | **1.229** | **<0.001** |
| Limonene-Control | **1.544** | **<0.001** | **1.409** | **<0.001** | **0.881** | **0.024** | 0.940 | 0.810 |
| MBOA-Control | **1.685** | **<0.001** | 1.058 | 0.726 | **0.544** | **<0.001** | 0.968 | 0.967 |
| MHPA-Control | **1.670** | **<0.001** | 1.108 | 0.117 | 0.973 | 0.983 | 0.960 | 0.963 |
| Mix-Control | **1.693** | **<0.001** | **1.305** | **<0.001** | **0.627** | **<0.001** | 1.009 | 1.000 |
| Limonene-DMPP | **1.182** | **0.001** | 0.929 | 0.452 | **0.876** | **0.015** | **0.765** | **<0.001** |
| MBOA-DMPP | **1.289** | **<0.001** | **0.698** | **<0.001** | **0.541** | **<0.001** | **0.788** | **<0.001** |
| MHPA-DMPP | **1.277** | **<0.001** | **0.731** | **<0.001** | 0.967 | 0.960 | **0.781** | **<0.001** |
| Mix-DMPP | **1.295** | **<0.001** | **0.861** | **0.004** | **0.624** | **<0.001** | **0.821** | **0.028** |
| MBOA-Limonene | 1.091 | 0.263 | **0.751** | **<0.001** | **0.618** | **<0.001** | 1.030 | 0.991 |
| MHPA-Limonene | 1.081 | 0.386 | **0.787** | **<0.001** | 1.104 | 0.143 | 1.021 | 0.999 |
| Mix-Limonene | 1.096 | 0.208 | 0.926 | 0.407 | **0.712** | **<0.001** | 1.074 | 0.911 |
| MHPA-MBOA | 0.991 | 1.000 | 1.047 | 0.859 | **1.788** | **<0.001** | 0.992 | 1.000 |
| Mix-MBOA | 1.005 | 1.000 | **1.233** | **<0.001** | **1.153** | **0.017** | 1.042 | 0.987 |
| Mix-MHPA | 1.014 | 0.999 | **1.178** | **0.001** | **0.645** | **<0.001** | 1.051 | 0.980 |

**B.**

| Comparison | Day 0 | | Day 5 | | Day 9 | | Day 14 | |
|---|---|---|---|---|---|---|---|---|
| | ratio | p adj | ratio | p adj | ratio | p adj | ratio | p adj |
| DMPP-Control | **1.247** | **<0.001** | **1.358** | **<0.001** | 1.000 | 1.000 | **1.259** | **<0.001** |
| Limonene-Control | **1.588** | **<0.001** | 1.070 | 0.626 | **0.654** | **<0.001** | 0.972 | 0.980 |
| MBOA-Control | **1.642** | **<0.001** | 1.048 | 0.853 | **0.572** | **<0.001** | 0.992 | 1.000 |
| MHPA-Control | **1.626** | **<0.001** | 1.060 | 0.701 | 0.981 | 0.997 | 1.061 | 0.680 |
| Mix-Control | **1.690** | **<0.001** | 1.081 | 0.383 | **0.622** | **<0.001** | 1.000 | 1.000 |
| Limonene-DMPP | **1.274** | **<0.001** | **0.788** | **<0.001** | **0.654** | **<0.001** | **0.772** | **<0.001** |
| MBOA-DMPP | **1.317** | **<0.001** | **0.772** | **<0.001** | **0.572** | **<0.001** | **0.788** | **<0.001** |
| MHPA-DMPP | **1.304** | **<0.001** | **0.780** | **<0.001** | 0.981 | 0.997 | **0.843** | **0.001** |
| Mix-DMPP | **1.355** | **<0.001** | **0.796** | **<0.001** | **0.622** | **<0.001** | **0.794** | **<0.001** |
| MBOA-Limonene | 1.034 | 0.962 | 0.979 | 0.997 | **0.875** | **0.014** | 1.021 | 0.996 |
| MHPA-Limonene | 1.024 | 0.994 | 0.990 | 1.000 | **1.499** | **<0.001** | 1.092 | 0.250 |

| | | | | | | | | |
|---|---|---|---|---|---|---|---|---|
| Mix-Limonene | 1.064 | 0.641 | 1.010 | 1.000 | 0.951 | 0.855 | 1.029 | 0.993 |
| MHPA-MBOA | 0.990 | 1.000 | 1.011 | 1.000 | **1.714** | **<0.001** | 1.070 | 0.547 |
| Mix-MBOA | 1.029 | 0.980 | 1.032 | 0.971 | 1.087 | 0.394 | 1.008 | 1.000 |
| Mix-MHPA | 1.039 | 0.951 | 1.020 | 0.996 | **0.634** | **<0.001** | 0.942 | 0.829 |

**Table S18:** Post hoc test results for differences in $NO_3^-$ as a fraction of applied N among treatments within the **A.** planted and **B.** bare-soil compartments. Pairwise comparisons of estimated marginal means (EMMs) from a gamma–log generalized linear model (GLM) were performed. Significant differences relative to the control and between treatments are shown in bold.

**A.**

| Comparison | Day 0 | | Day 5 | | Day 9 | | Day 14 | |
|---|---|---|---|---|---|---|---|---|
| | ratio | p adj | ratio | p adj | ratio | p adj | ratio | p adj |
| DMPP-Control | 0.913 | 0.929 | 0.972 | 1.000 | 0.967 | 0.999 | 1.073 | 0.976 |
| Limonene-Control | 1.045 | 0.997 | 1.267 | 0.154 | **1.430** | **0.003** | 1.100 | 0.937 |
| MBOA-Control | 1.262 | 0.139 | **1.876** | **<0.001** | **1.577** | **<0.001** | 1.186 | 0.465 |
| MHPA-Control | 1.089 | 0.951 | 1.148 | 0.686 | 1.043 | 0.998 | 0.981 | 1.000 |
| Mix-Control | **1.522** | **<0.001** | **2.391** | **<0.001** | **2.597** | **<0.001** | **2.514** | **<0.001** |
| Limonene-DMPP | 1.144 | 0.710 | 1.304 | 0.077 | **1.479** | **0.001** | 1.025 | 1.000 |
| MBOA-DMPP | **1.382** | **0.010** | **1.931** | **<0.001** | **1.632** | **<0.001** | 1.105 | 0.896 |
| MHPA-DMPP | 1.193 | 0.465 | 1.182 | 0.486 | 1.079 | 0.966 | 0.914 | 0.951 |
| Mix-DMPP | **1.667** | **<0.001** | **2.461** | **<0.001** | **2.687** | **<0.001** | **2.344** | **<0.001** |
| MBOA-Limonene | 1.208 | 0.344 | **1.481** | **0.001** | 1.103 | 0.903 | 1.078 | 0.977 |
| MHPA-Limonene | 1.043 | 0.998 | 0.906 | 0.914 | **0.729** | **0.013** | 0.892 | 0.899 |
| Mix-Limonene | **1.457** | **0.001** | **1.887** | **<0.001** | **1.816** | **<0.001** | **2.286** | **<0.001** |
| MHPA-MBOA | 0.863 | 0.659 | **0.612** | **<0.001** | **0.661** | **<0.001** | 0.827 | 0.431 |
| Mix-MBOA | 1.206 | 0.354 | 1.274 | 0.110 | **1.646** | **<0.001** | **2.121** | **<0.001** |
| Mix-MHPA | **1.398** | **0.010** | **2.083** | **<0.001** | **2.490** | **<0.001** | **2.563** | **<0.001** |

**B.**

| Comparison | Day 0 | | Day 5 | | Day 9 | | Day 14 | |
|---|---|---|---|---|---|---|---|---|
| | ratio | p adj | ratio | p adj | ratio | p adj | ratio | p adj |
| DMPP-Control | 0.913 | 0.929 | 0.972 | 1.000 | 0.967 | 0.999 | 1.073 | 0.976 |
| Limonene-Control | 1.045 | 0.997 | 1.267 | 0.154 | **1.430** | **0.003** | 1.100 | 0.937 |
| MBOA-Control | 1.262 | 0.139 | **1.876** | **<0.001** | **1.577** | **<0.001** | 1.186 | 0.465 |
| MHPA-Control | 1.089 | 0.951 | 1.148 | 0.686 | 1.043 | 0.998 | 0.981 | 1.000 |
| Mix-Control | **1.522** | **<0.001** | **2.391** | **<0.001** | **2.597** | **<0.001** | **2.514** | **<0.001** |
| Limonene-DMPP | 1.144 | 0.710 | 1.304 | 0.077 | **1.479** | **0.001** | 1.025 | 1.000 |
| MBOA-DMPP | **1.382** | **0.010** | **1.931** | **<0.001** | **1.632** | **<0.001** | 1.105 | 0.896 |
| MHPA-DMPP | 1.193 | 0.465 | 1.182 | 0.486 | 1.079 | 0.966 | 0.914 | 0.951 |
| Mix-DMPP | **1.667** | **<0.001** | **2.461** | **<0.001** | **2.687** | **<0.001** | **2.344** | **<0.001** |
| MBOA-Limonene | 1.208 | 0.344 | **1.481** | **0.001** | 1.103 | 0.903 | 1.078 | 0.977 |
| MHPA-Limonene | 1.043 | 0.998 | 0.906 | 0.914 | **0.729** | **0.013** | 0.892 | 0.899 |
| Mix-Limonene | **1.457** | **0.001** | **1.887** | **<0.001** | **1.816** | **<0.001** | **2.286** | **<0.001** |

| | | | | | | | | |
|---|---|---|---|---|---|---|---|---|
| MHPA-MBOA | 0.863 | 0.659 | **0.612** | **<0.001** | **0.661** | **<0.001** | 0.827 | 0.431 |
| Mix-MBOA | 1.206 | 0.354 | 1.274 | 0.110 | **1.646** | **<0.001** | **2.121** | **<0.001** |
| Mix-MHPA | **1.398** | **0.010** | **2.083** | **<0.001** | **2.490** | **<0.001** | **2.563** | **<0.001** |
